# Profiling ubiquitin signaling with UBIMAX reveals DNA damage- and SCF^β^^TRCP^-dependent ubiquitylation of the actin-organizing protein Dbn1

**DOI:** 10.1101/2023.05.15.540799

**Authors:** Camilla S. Colding-Christensen, Ellen S. Kakulidis, Javier Arroyo-Gomez, Ivo A. Hendriks, Connor Arkinson, Zita Fábián, Agnieszka Gambus, Niels Mailand, Julien P. Duxin, Michael L. Nielsen

**Author notes:** These authors contributed equally to this work (Camilla S. Colding-Christensen, Ellen S. Kakulidis). Correspondence and requests for materials should be addressed to Julien P. Duxin and Michael L. Nielsen.

## Abstract

Ubiquitin widely modifies proteins, thereby regulating most cellular functions. The complexity of ubiquitin signalling necessitates unbiased methods enabling global detection of dynamic protein ubiquitylation. Here, we describe UBIMAX (<u>UB</u>iquitin target Identification by <u>M</u>ass spectrometry in *<u>X</u>enopus* egg extracts), which enriches ubiquitin-conjugated proteins and quantifies regulation of protein ubiquitylation under precise and adaptable conditions. We benchmark UBIMAX by investigating DNA double-strand break-responsive ubiquitylation events, identifying previously known targets and revealing the actin-organising protein Dbn1 as a novel major target of DNA damage-induced ubiquitylation. We find that Dbn1 is targeted for proteasomal degradation by the SCF^β-Trcp1^ ubiquitin ligase, in a conserved mechanism driven by ATM-mediated phosphorylation of a previously uncharacterized β-Trcp1 degron containing an SQ motif. We further show that this degron is sufficient to induce DNA-damage dependent protein degradation of a model substrate. Collectively, we demonstrate UBIMAX’s ability to identify novel targets of stimulus-regulated ubiquitylation and reveal an SCF^β-Trcp1^-mediated ubiquitylation mechanism controlled directly by the apical DNA damage response kinases.

## Introduction

Ubiquitin is a small 76 amino acid protein, which can be attached via its C-terminus to target proteins via the catalysis by specific ubiquitin conjugating enzymes (Ciehanover et al., 1978; Goldknopf and Busch, 1977; Pickart, 2001). It is estimated that this dynamic post-translational modification (PTM), ubiquitylation, regulates nearly all cellular functions (Swatek and Komander, 2016). As ubiquitin can be attached to target proteins as monomers and as chains of different topologies, the resulting signal can be highly complex. These signals are decoded by proteins with the ability to interact with specific ubiquitin topologies. Such proteins are often effectors of cellular pathways, allowing ubiquitylation topologies to regulate distinct cellular functions (Swatek and Komander, 2016; Yau and Rape, 2016). For instance, many aspects of the DNA damage response (DDR) are regulated by K63- and K48-ubiquitin signalling, including recruitment of effectors to the DNA damage site and choice of repair pathway (Schwertman et al., 2016). One example is the DNA double-strand break (DSB)-induced K48-linked ubiquitylation of the Ku complex, which causes its eviction from DNA, thus regulating the DSB repair process (Feng and Chen, 2012; Ismail et al., 2015; Postow et al., 2008). While extensive biochemical and molecular analyses have been required for such investigations, the breadth and complexity of ubiquitin signalling has prompted the need for unbiased and global ubiquitin detection methods.

Mass spectrometry (MS)-based proteomics has emerged as a valuable tool for studying the ubiquitin landscape and has been widely used to investigate ubiquitylation events in the DDR and DNA repair processes (Foster et al., 2021). Over the last decade, numerous MS-based approaches have been established to identify ubiquitylated proteins, determine the amino acid acceptor sites, establish the chain-topology of ubiquitylation, and identify the enzymes involved (Foster et al., 2021; Sun and Zhang, 2022; Trulsson and Vertegaal, 2022). Particularly the methods for identifying ubiquitylation sites have been extensively used for profiling ubiquitylation responses (Akimov et al., 2018a; Akimov et al., 2018b; Kim et al., 2011; Wagner et al., 2011; Xu et al., 2010). Although these methods have provided valuable insights into the ubiquitin landscape, they are limited in providing quantitative information of the ubiquitylated target. While methods for identification of ubiquitylation on the protein level can provide such quantitative information about ubiquitylated proteoforms (Akimov et al., 2014; Danielsen et al., 2011; Hjerpe et al., 2009; Lopitz-Otsoa et al., 2012; Peng et al., 2003), these methods are often challenged by a lack of specificity in enriching ubiquitin-conjugated *versus*-interacting proteins. Furthermore, current methods have limitations when it comes to capturing steady-state systems and targeting specific events in response to a particular stimulus. For instance, it is challenging to generate site-specific DNA lesions in cellular model systems, prompting the need for developing an *in vitro* system to precisely study ubiquitylation responses to defined stimuli.

The *Xenopus* egg extract model system has been extensively used for biochemical and molecular studies of DNA metabolism and genome maintenance mechanisms (Hoogenboom et al., 2017; Raspelli et al., 2017; Sannino et al., 2016). This is chiefly owing to the possibility of investigating key biological processes with high spatiotemporal resolution in the absence of essential proteins, and upon the ready addition of recombinant proteins or specific inhibitors. Recently, *Xenopus* egg extracts have been coupled to MS-based proteomic analyses to study quantifiable changes in protein recruitment to damaged DNA (Larsen et al., 2019; Raschle et al., 2015).

Here we describe a new MS-based method, referred to as UBIMAX (<u>UB</u>iquitin target Identification by <u>M</u>ass spectrometry in *<u>X</u>enopus* egg extracts), which allows for detection of global, specific, and quantifiable changes in *de novo* protein ubiquitylation under precise and modifiable biological conditions in *Xenopus* egg extracts. As proof of principle, we use UBIMAX to identify previously characterized DNA damage-induced ubiquitylation events alongside several novel targets of DNA damage-specific ubiquitylation in response to DSBs and DNA-protein crosslinks (DPCs). From this we discovered that the actin-organizing protein Dbn1, which has not been previously linked to the DDR, is a novel prominent ubiquitylation target in response to DSBs. Using *Xenopus* egg extract and human cells, we demonstrate that DSB-induced Dbn1 ubiquitylation depends on the apical DDR kinase ATM, is mediated by the Skp1-Cul1-F-box^β-Trcp1^ (SCF^β-Trcp1^) complex, and leads to proteasomal degradation of Dbn1. Additionally, we uncover a DNA damage-responsive β-Trcp1-degron in the C-terminal unstructured region of Dbn1 and show that it is necessary and sufficient for conferring DSB-induced ubiquitylation and degradation of a target protein. Collectively, we demonstrate the capability of UBIMAX to identify novel players in ubiquitylation responses under specific biological conditions of interest. Furthermore, by deciphering the mechanism of DSB-induced Dbn1 ubiquitylation identified by UBIMAX, we reveal a variant β-Trcp1 degron that mediates a DDR-and SCF^β-Trcp1^-specific ubiquitylation and degradation programme.

## Results

### UBIMAX efficiently and specifically detects ubiquitin-conjugated proteins

To establish a method for identification of endogenous ubiquitylation events in response to specific stimuli, we combined high-resolution mass spectrometry (MS) with the malleable *Xenopus* egg extract system (Raspelli et al., 2017). Briefly, we supplemented high speed supernatant interphase egg extracts (HSS) with recombinant 6xHis-tagged ubiquitin prior to inducing a stimulus-driven response (Figure 1A). Following a specific stimulation, endogenous and supplemented ubiquitin was allowed time to conjugate onto target proteins, enabling highly stringent enrichment of target proteins via pulldown of His-ubiquitin. Next, the enriched ubiquitylated proteins were digested on-beads using trypsin, with the resulting peptides purified and concentrated on C_18_-StageTips (Rappsilber et al., 2003) and followed by their characterization via label-free quantitative MS (Cox et al., 2014).

**Figure 1.**
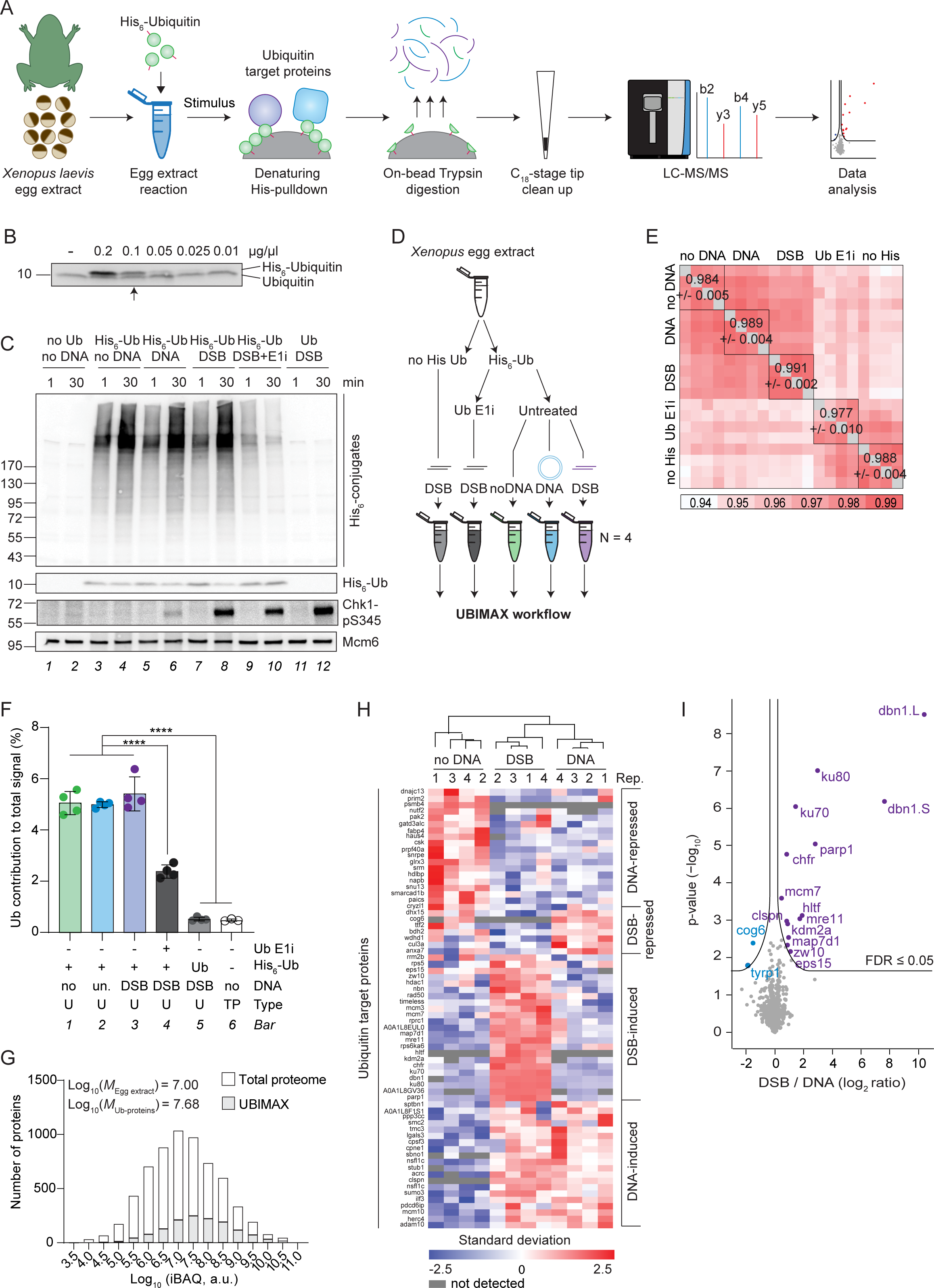
UBIMAX efficiently and specifically detects ubiquitin-conjugated proteins in response to DSBs. **A.** Schematic representation of the experimental system and workflow for UBIMAX. LC-MS/MS; liquid chromatography-tandem mass spectrometry **B.** Recombinant His_6_-ubiquitin was added to egg extracts at the indicated final concentrations and samples immediately transferred to sample buffer for WB against ubiquitin. The arrow indicates the final concentration of His_6_-ubiquitin used in all subsequent experiments, 0.1 µg/µl, unless otherwise stated. **C-D.** Egg extracts were supplemented with DMSO or ubiquitin E1 inhibitor (“E1i”) prior to addition of recombinant untagged ubiquitin (“Ub”) or His_6_-Ubiquitin (“His_6_-Ub”) as indicated. Reactions were initiated by addition of buffer (“no DNA”), undamaged plasmid DNA (“DNA”) or linearized plasmid DNA (“DSB”). For WB analysis (**C**), samples were transferred to sample buffer at 1 or 30 min after initiation of the reaction and blotted against the His-tag or Chk1-pS345. Mcm6 served as a loading control. For UBIMAX (experimental outline in **D**), reactions were performed in quadruplicate from the same batch of egg extracts. Samples were transferred to denaturing pulldown buffer 30 min after initiation of the reaction and subjected to the UBIMAX workflow as outlined in (A). no His Ub, untagged recombinant ubiquitin; Ub E1i, ubiquitin E1 inhibitor. **E.** Pearson correlation matrix of the experiment outlined in Figure 1D. Within-replicate average and ±standard deviation of Pearson correlation is indicated. **F.** Mean percent contribution of ubiquitin peptide abundance to summed peptide intensity across four replicates of the UBIMAX samples (“U”) as well as for three replicates of a total egg extract proteome (“TP”) derived from an independent experiment (data not shown). Error bars represent standard deviations. Significance was determined by one-way ANOVA with Tukey’s multiple comparisons test for all pairwise comparisons. no, no DNA (buffer); un., undamaged plasmid DNA. **G.** Depth of sequencing illustrated as distribution of Log_10_(iBAQ)-values of the ubiquitylated proteins detected by UBIMAX compared to the proteins detected in a total egg extract proteome derived from an independent experiment (data not shown). Frequency distribution medians (*“M*”) are shown at the top left corner of the graph. iBAQ, intensity-based absolute quantification; a.u., arbitrary units. **H.** Hierarchical clustering analysis of Z-scored abundances of the ubiquitylated proteins robustly changing with DNA treatment. Gene names are provided on the left. If gene names could not be annotated, the UniProt ID is given instead. Rep., replicate. **I.** Volcano plot analysis comparing ubiquitylated proteins enriched from DSB versus undamaged DNA-treated samples. Purple and blue dots indicate significantly enriched and depleted ubiquitylated proteins, respectively. Significance was determined by two-tailed Student’s *t* test, with permutation-based FDR-control with S0 = 0.1 and 2500 rounds of randomization, to ensure FDR ≤ 0.05. N=4.

To minimize non-specific ubiquitylation events, we ensured that recombinant 6xHis-tagged ubiquitin was titrated to equimolar amounts of endogenous ubiquitin (Figure 1B). Under these conditions, recombinant 6xHis-tagged ubiquitin was efficiently conjugated onto target proteins (Figures 1C lanes 3-8 and S1A), thus allowing for efficient enrichment of these proteins using the UBIMAX approach.

As ubiquitylation constitutes an important part of the response to DNA damage (Schwertman et al., 2016), we chose DNA double-strand breaks (DSBs) as our stimulus of choice for benchmarking UBIMAX. To elicit a DSB response, we added linearized plasmid DNA to egg extracts, while omission of DNA or addition of circular undamaged plasmid DNA served as controls. Importantly, addition of recombinant 6xHis-tagged ubiquitin did not activate the DNA damage response (DDR) in the absence of DNA (Figure 1C, lanes 3-4), nor did it affect DDR activation by the linearized plasmid, as indicated by Chk1-S345 phosphorylation (Figure 1C, lane 8). DSB repair was also unaffected by addition of recombinant 6xHis-tagged ubiquitin, and linearized plasmids were ligated by non-homologous end joining (NHEJ) irrespectively of the presence or absence of recombinant ubiquitin (Di Virgilio and Gautier, 2005; Graham et al., 2016; Pfeiffer and Vielmetter, 1988) (Figure S1B and quantifications in Figure S1C). In contrast, NHEJ-mediated repair of DSBs was partially impaired in the presence of a ubiquitin E1 inhibitor (Gallina et al., 2021), confirming the relevance of *de novo* ubiquitylation for DSB repair in egg extracts (Figure S1B-C). Collectively, we established appropriate conditions to study protein ubiquitylation in response to DSBs.

Next, we sought to profile DSB-induced ubiquitylation events using UBIMAX. To ensure that the method enriches ubiquitin-conjugated target proteins, as opposed to ubiquitin-interacting proteins, we performed all enrichments under denaturing conditions. To distinguish between specific and non-specific enrichment of proteins, we additionally performed reactions in egg extracts supplemented with either recombinant 6xHis-tagged or untagged ubiquitin (Figure 1D). As an additional control, we performed reactions in the presence or absence of ubiquitin E1 inhibitor to block ubiquitylation. Finally, as described above, we investigated DSB-induced ubiquitylation events by adding linearized plasmid DNA, circular plasmid DNA, or no plasmid DNA to individual reactions. We performed all reactions in quadruplicate and collected samples 30 minutes after addition of DNA. Samples were subsequently subjected to the UBIMAX workflow (Figure 1A). Overall, we observed very high reproducibility across all replicates and sample groups (*R* = 0.94 – 0.99) (Figure 1E), supporting that the dynamics of the investigated ubiquitylation landscape were highly specific. This was corroborated by principal component analysis (PCA), which revealed a large variation between controls and ubiquitin target enriched sample groups (Figure S1D). Correspondingly, UBIMAX robustly separated the ubiquitin target enriched sample groups based on the introduced DNA stimuli (Figure S1E).

To assess the efficiency of our enrichment approach, we analysed the average contribution of ubiquitin peptides to total sample signal (Figure S1F) and found it significantly higher in ubiquitin target enriched sample groups compared to each of the controls (Figure 1F, compare bars 1-3 with 4 and 5). Importantly, the ubiquitin contribution in the untagged ubiquitin control was similar to that of a total proteome (Figure 1F, compare bars 5 and 6). This shows that our denaturing His-ubiquitin enrichment approach is highly efficient with approximately 90% of the ubiquitin signal being specific (Figure 1F, ratio of bars 1-3 and 5). Furthermore, ubiquitin contribution in the ubiquitin target enriched sample groups was approximately two-fold higher than in the ubiquitin E1 inhibitor control (Figure 1F, compare bars 1-3 with 4). As a large fraction of recombinant 6xHis-tagged ubiquitin is left unconjugated in the presence of the ubiquitin E1 inhibitor (Figure 1C, lanes 9 and 10), this indicates that while non-conjugated His-ubiquitin is efficiently recovered in the presence of ubiquitin E1 inhibitor, a similar amount of endogenous ubiquitin is additionally recovered in the stimulus groups. This observation is presumably due to the equal concentration of endogenous and recombinant 6xHis-tagged ubiquitin (Figure 1B) and suggest that these are equally conjugated onto target proteins.

Finally, as ubiquitylation is a sub-stoichiometric PTM, we assessed the abundance bias of UBIMAX. Overall, we observed a large dynamic range (∼7000-fold) with a distribution of ubiquitylated proteins identified by UBIMAX similar to that of a total egg extract proteome (Log_10_ *M* = 7.68 and 7.00, respectively) (Figure 1G), indicating a relatively modest abundance bias for UBIMAX. Collectively, these data show that UBIMAX is an efficient and robust method for identifying specific ubiquitylation events in *Xenopus* egg extracts.

### UBIMAX identifies DSB-induced protein ubiquitylation

From quadruplicate experiments, UBIMAX identified 786 significantly His-pulldown enriched and *de novo* ubiquitylated proteins across the ubiquitin target enriched sample groups (Figure S1G, left). To elucidate subsets of proteins whose ubiquitylation status were consistently up-or downregulated across replicates, we interrogated these 786 proteins further and identified four clusters of specifically regulated ubiquitylated proteins in response to the DNA treatments (Figure 1H). Gene Ontology (GO) term enrichment analysis of individual clusters revealed that addition of exogenous DNA to *Xenopus* egg extracts induced ubiquitylation of proteins involved in e.g. immune and inflammatory processes, whereas ubiquitylation of proteins involved in DNA repair, DNA replication, and checkpoint control was upregulated in response to DSBs (Figures 1H and S1H). This latter group included well-known DDR proteins such as the Ku70-Ku80 dimer, the Mre11-Rad50-Nbs1 (MRN) complex, and Parp1 (Figure S1I). It also included DNA replication factors such as Mcm3, Mcm7, and Timeless.

Next, we interrogated the 39 proteins which showed a significant regulation of ubiquitylation status upon stimulation with either undamaged or DSB-containing plasmid DNA (S1G, right). Volcano plot analysis of these ubiquitylation events confirmed the previously described DSB-induced ubiquitylation of Ku80, Parp1, Mre11, and Claspin (Jachimowicz et al., 2019; Kim et al., 2018; Liu et al., 2013; Mailand et al., 2006; Peschiaroli et al., 2006; Postow and Funabiki, 2013; Postow et al., 2008) (Figure 1I). Moreover, we also detected enrichment of ubiquitylated Hltf and Chfr, two ubiquitin E3 ligases known to auto-ubiquitylate upon DNA damage (Chaturvedi et al., 2002; Lin et al., 2011; Liu et al., 2013). Finally, we detected DSB-induced ubiquitylation of proteins not previously described as being modified upon DSBs, including Mcm7. Strikingly, the most prominently induced ubiquitylated protein detected in response to DSBs was the actin-organizing protein Dbn1, a protein not previously connected with the DSB response. In conclusion, we demonstrate the ability of UBIMAX to reveal regulation of protein ubiquitylation events in response to DNA damage.

### UBIMAX identifies DNA damage specific ubiquitylation events

To further assess the capability of UBIMAX to detect ubiquitylation events triggered by a specific stimulus, we used UBIMAX to analyse the ubiquitylation response to different DNA lesions. To this end, we used plasmids carrying either the previously described *Haemophilus parainfluenzae* methyltransferase M.HpaII crosslinked at a single-stranded DNA gap (“ssDNA-DPC”) (Larsen et al., 2019), or the *Saccharomyces cerevisiae* recombinase Flp crosslinked at a single-strand break (“SSB-DPC”) (Nielsen et al., 2009) (Figure 2A). Repair of these DPC substrates have previously been shown to require ubiquitylation (Duxin et al., 2014; Larsen et al., 2019; Serbyn et al., 2021), thus making these DNA lesions relevant for UBIMAX analysis. Furthermore, while both are DPC lesions, the nature of the protein adduct and the DNA context (located on ssDNA *versus* at a SSB) are different, thus serving as a suitable test for the specificity of UBIMAX in distinguishing these responses.

**Figure 2.**
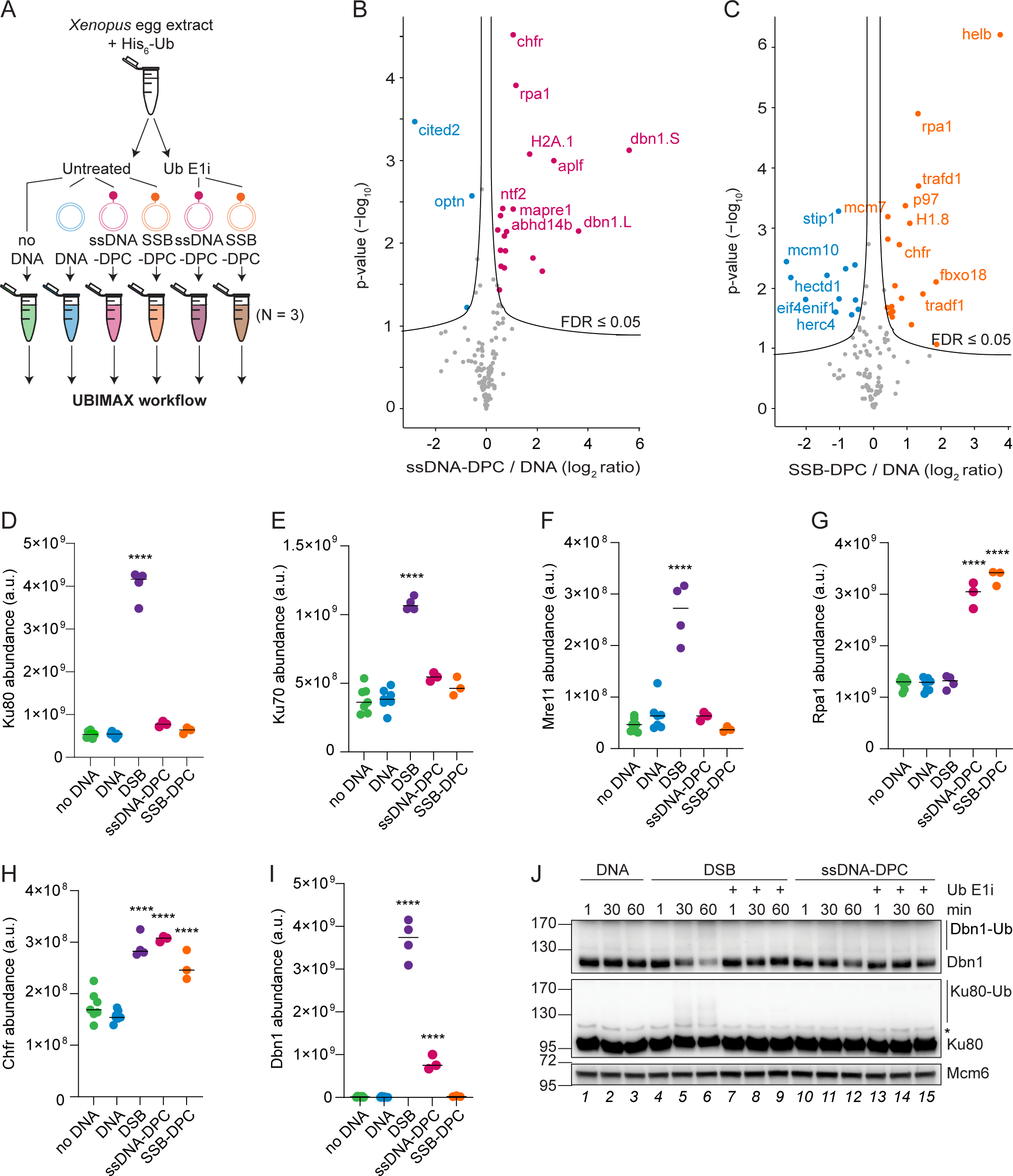
UBIMAX identifies DNA damage specific ubiquitylation events and detects DSB-induced ubiquitylation of Dbn1. **A.** Experimental outline for the UBIMAX experiment profiling ubiquitylated proteins in response to DPC-containing substrates. Egg extracts were left untreated or supplemented with ubiquitin E1 inhibitor (“Ub E1i”) prior to addition of His_6_-Ubiquitin (“His_6_-Ub”). Reactions were initiated by addition of buffer (“no DNA”), undamaged plasmid DNA (“DNA”), plasmids carrying the M.HpaII protein crosslinked at a single-stranded DNA gap (“ssDNA-DPC”), or plasmids carrying the Flp protein crosslinked at a single-strand break (“SSB-DPC”). Reactions were performed in triplicate from the same batch of egg extracts. Samples were transferred to denaturing pulldown buffer 30 min after initiation of the reaction and subjected to the UBIMAX workflow as outlined in Figure 1A. **B-C.** Volcano plot analysis comparing ubiquitylated proteins enriched from ssDNA-DPC (**B**) or SSB-DPC (**C**) versus DNA-treated samples. Pink/orange and blue dots indicate significantly enriched and depleted ubiquitylated proteins, respectively. Significance was determined by two-tailed Student’s *t* test, with permutation-based FDR-control with S0 = 0.1 and 2500 rounds of randomization, to ensure an FDR ≤ ss0.05. Ubiquitylated proteins with FDR ≤ 0.01 are labelled. N=3. **D-I.** Abundance distributions of Ku80 (**D**), Ku70 (**E**), Mre11 (**F**), Rpa1 (**G**), Chfr (**H**) and Dbn1 (**I**) across the ubiquitin target enriched samples of the UBIMAX experiments profiling protein ubiquitylation in response to DSBs (Figure 1D) and DPCs (A), respectively. Horizontal lines indicate the median and significance was determined by one-way ANOVA with Dunnett’s multiple comparisons test for all conditions against undamaged DNA with a cut-off of p-value ≤ 0.01. N=3-7. a.u., arbitrary units. **J.** Egg extracts were left untreated or supplemented with ubiquitin E1 inhibitor prior to initiation of the reactions by addition of either undamaged plasmid DNA (“DNA”), linearized plasmid DNA (“DSB”), or plasmids carrying a DPC at a ssDNA gap (“ssDNA-DPC”). Samples were transferred to sample buffer at the indicated times and analysed by WB using antibodies against Dbn1 and Ku80. Mcm6 served as a loading control. * denotes an unspecific band.

For profiling ubiquitylation events by UBIMAX in response to each of the DPC-containing plasmids, we also included the ubiquitin E1 inhibitor control. Reactions were performed in triplicate with samples collected 30 min after addition of DNA and subjected to the UBIMAX workflow (Figure 1A; Figure S2A). From this, we detected distinct *de novo* protein ubiquitylation events induced by either ssDNA-DPC or SSB-DPC (Figures 2B-C). While some proteins were found ubiquitylated in response to both substrates (e.g. Chfr, and Rpa1), UBIMAX also detected proteins uniquely ubiquitylated in response to either plasmid (e.g. Aplf for ssDNA-DPC; HelB for SSB-DPC). Next, we compared the proteins showing upregulation of ubiquitylation in response to the different DNA lesions (DSB vs ssDNA-DPC vs SSB-DPC) as detected by UBIMAX (Figure S2B). From this, we found that each type of DNA damage predominantly induced DNA damage-specific ubiquitylation events, with a few factors ubiquitylated in more than one condition. Consistent with their role in NHEJ and DSB repair, Ku80, Ku70, and Mre11 were specifically ubiquitylated in the presence of the DSB containing plasmid (Figures 2D-F). In contrast, ubiquitylation of the ssDNA binding protein RPA (Elia et al., 2015) was greatly stimulated by the DPC lesions flanked by either ssDNA or a SSB, consistent with RPA binding to these substrates (in the case of the SSB-DPC, presumably once the SSB has been resected) (Figure 2G). The only protein ubiquitylated in response to all three DNA lesions was Chfr (Figure 2H), which is recruited to DNA damage sites in a poly(ADP-ribose) (PAR)-dependent manner (Liu et al., 2013), with PARylation likely occurring at all of these DNA lesions. Finally, we found that the actin-organizing protein Dbn1 was ubiquitylated in response to DSBs and, to a lesser degree, DPCs flanked by a ssDNA gap but not in response to DPCs flanked by a SSB (Figure 2I). Consistent with the UBIMAX data, western blot (WB) analysis confirmed that Dbn1 and Ku80 were ubiquitylated primarily in response to DSBs and to a much lesser extent in response to ssDNA-DPCs (Figure 2J, lanes 4-6 and 10-12). In conclusion, we demonstrate the precision of UBIMAX in detecting DNA damage specific *de novo* protein ubiquitylation events and identify the DNA damage-induced ubiquitylation of the actin-organizing protein Dbn1 primarily in response to DSBs.

### DDR-dependent ubiquitylation of Dbn1 results in proteasomal degradation

To further validate the ability of UBIMAX to identify novel ubiquitylated substrates, we sought to characterize the previously unknown, damage-induced ubiquitylation of Dbn1. To this end, we first generated a Dbn1 antibody that efficiently immunodepleted Dbn1 from egg extracts (Figure S3A). We then performed denaturing His-ubiquitin pulldowns from mock-or Dbn1 immunodepleted egg extracts supplemented with 6xHis-tagged ubiquitin and DSB plasmid and analysed the recovered proteins by WB (Figure 3A). The Dbn1 signal recovered from mock-immunodepleted samples migrated as a smear 30 min after addition of DSBs and peaked at 60 min (Figure 3A, lanes 1-3). In contrast, no signal was detected in Dbn1-immunodepleted extracts (Figure 3A, lanes 4-6), validating the prominent poly-ubiquitylation of Dbn1 following DSBs.

**Figure 3.**
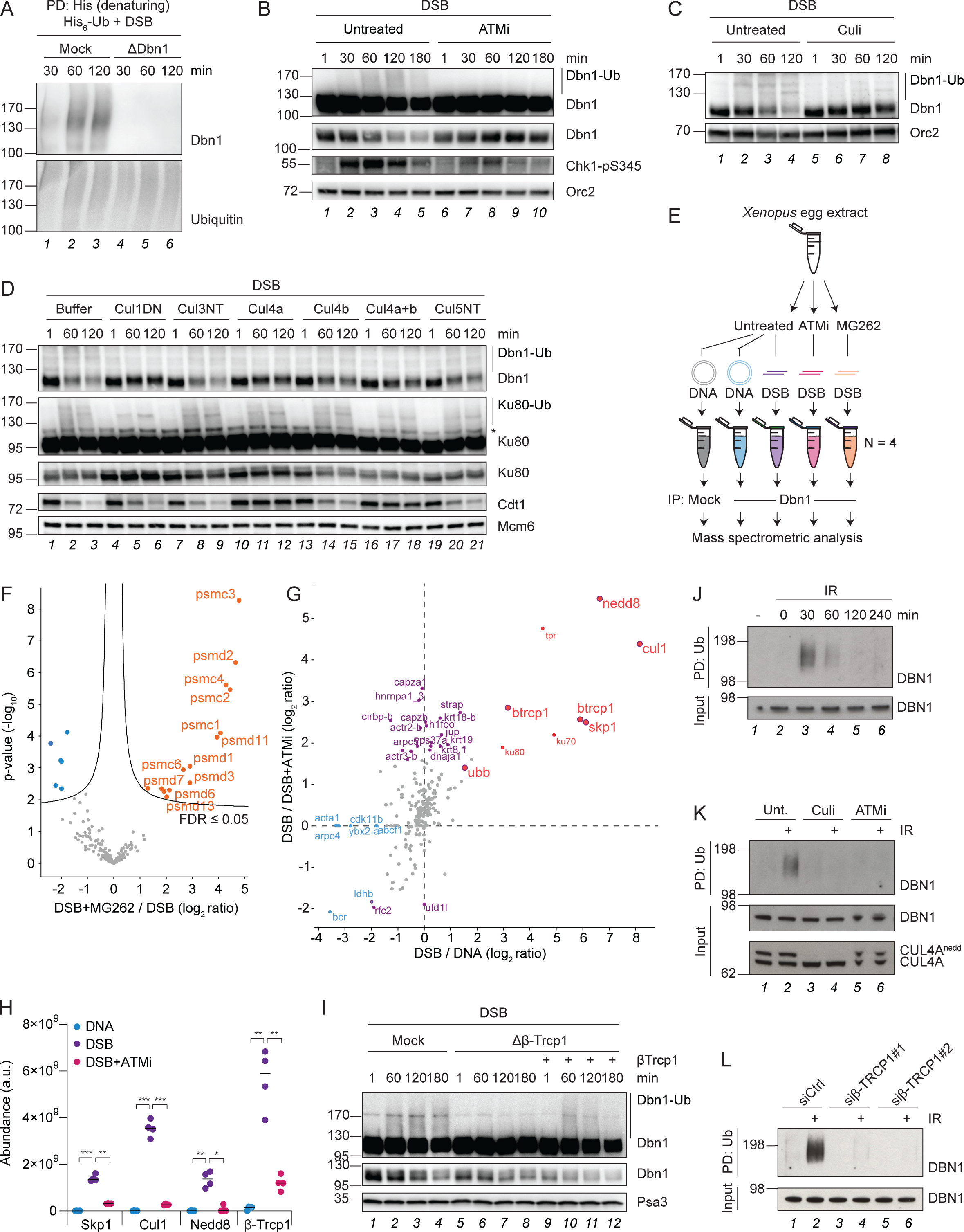
DSB-induced ubiquitylation of Dbn1 depends on ATM and is mediated by the SCF^β-Trcp1^ E3 ligase. **A.** Mock- or Dbn1-immunodepleted egg extracts (Figure S3A) were supplemented with His_6_-ubiquitin (“His_6_-Ub”) prior to addition of linearized plasmid DNA (“DSB”). Samples were collected at the indicated times and subjected to denaturing His-ubiquitin pulldown followed by WB analysis using antibodies against Dbn1. Ubiquitin served as a pulldown control. Immunodepletion control is provided in Figure S3A. PD, pulldown. **B.** Egg extracts were left untreated or supplemented with ATM inhibitor (“ATMi”) prior to addition of linearized plasmid DNA. Samples were transferred to sample buffer at the indicated timepoints and analysed by WB using antibodies against Dbn1 (long and short exposures shown) and Chk1-pS345. Orc2 served as a loading control. **C.** Egg extracts were left untreated or supplemented with neddylation E1 inhibitor (“Culi”) prior to addition of linearized plasmid DNA. Samples were transferred to sample buffer at the indicated timepoints and analysed by WB using antibodies against Dbn1. Orc2 served as a loading control. **D.** Recombinant dominant negative Cullin proteins or buffer was mixed with egg extracts prior to addition of linearized plasmid DNA. Samples were transferred to sample buffer at the indicated timepoints and analysed by WB using antibodies against Dbn1, Ku80 (long and short exposures shown) and Cdt1. Mcm6 served as a loading control. * denotes an unspecific band. **E.** Experimental outline of Dbn1 IP-MS experiment. Egg extracts were left untreated or supplemented with ATM inhibitor or proteasome inhibitor (MG262) prior to addition of undamaged-(“DNA”) or linearized plasmid DNA (“DSB”) as indicated. Reactions were performed in quadruplicate from the same batch of egg extracts. Samples were collected at 60 min, subjected to mock-or Dbn1-immunoprecipitation as indicated and analysed by MS. **F.** Volcano plot analysis comparing the proteins enriched from Dbn1 IP-MS samples treated with DSB with versus without proteasomal inhibition. Orange and blue dots indicate significantly enriched and depleted ubiquitylated proteins, respectively. Significance was determined by two-tailed Student’s *t* test, with permutation-based FDR-control with S0 = 0.1 and 2500 rounds of randomization, to ensure an FDR ≤ 0.05. N=4. **G.** Scatter plot analysis of the Dbn1 IP-MS experiment detailed in (F). The mean difference in abundance between proteins enriched in linearized-and undamaged plasmid DNA-treated samples is plotted against that of linearized DNA without versus with ATM inhibition. Red and blue dots indicate proteins significantly enriched and depleted with Dbn1 immunoprecipitation in the presence of linearized plasmid DNA, respectively. Purple dots or outlines indicate proteins significantly changed in enrichment with Dbn1 immunoprecipitation upon ATM inhibition. Significance was determined by two-tailed Student’s *t* test with S0 = 0.1 and FDR ≤ 0.05. N=4. **H.** Abundance distributions of Skp1, Cul1, Nedd8, and β-Trcp1 across the indicated Dbn1 IP-MS conditions. Horizontal lines indicate the median and significance was determined by one-way ANOVA with Dunnett’s multiple comparisons test for all conditions shown against linearized plasmid DNA. N=4. a.u., arbitrary units. **I.** Recombinant β-Trcp1 protein or buffer was mixed with mock-or β-Trcp1-immunodepleted egg extracts as indicated prior to addition of linearized plasmid DNA. Samples were transferred to sample buffer at the indicated timepoints and analysed by WB using antibodies against Dbn1 (long and short exposures shown). Psa3 served as a loading control. **J.** HeLa cells were subjected to 10 Gy ionizing radiation (IR) and harvested after the indicated timepoints. Lysates were subjected to ubiquitin pulldown and analysed along with whole cell extracts (“input”) by WB using antibodies against DBN1. PD, pulldown; Ub, ubiquitin. **K.** HeLa cells were left untreated (“Unt.”) or treated with neddylation E1 inhibitor (“Culi”) or ATM inhibitor (“ATMi”) for 1 hour before being subjected or not to 10 Gy IR. Cells were harvested 30 minutes after irradiation and processed as described in (J). WB analysis of CUL4A served as a control for the neddylation E1 inhibitor. **L.** HeLa cells were transfected with control siRNA or two different siRNAs targeting β-Trcp1 for 72h before being subjected or not to 10 Gy IR. Cells were harvested 30 minutes after irradiation and processed as described in (J).

We next investigated the mechanism of DNA damage-induced Dbn1 ubiquitylation. We first addressed whether Dbn1 ubiquitylation required DDR activation. To investigate this, egg extract reactions were performed in the absence or presence of specific inhibitors of the apical DDR kinases, ATM and ATR, prior to stimulation with DSB plasmid DNA (Figures 3B and S3B). Addition of DSB plasmid DNA to egg extracts activated the DDR within minutes, as evidenced by the appearance of Chk1-S345 phosphorylation, followed by the appearance of Dbn1 ubiquitylation at 60 min (Figure 3B, lanes 1-5). Inhibition of either ATM or ATR inhibited DDR activation and DSB-induced Dbn1 ubiquitylation (Figures 3B lanes 6-10 and S3B lanes 7-18), suggesting that Dbn1 is ubiquitylated in response to DDR kinase activation.

To investigate the consequences of Dbn1 ubiquitylation, we next examined the major ubiquitin chain topologies present on the protein, as chain topology directs the functional consequences of protein ubiquitylation (Swatek and Komander, 2016; Yau and Rape, 2016). Particularly, K63- and K48-linked ubiquitin chains have been shown to orchestrate the response to DSBs (Schwertman et al., 2016). To this end, we assessed the extent of Dbn1 ubiquitylation in the presence of DSB plasmid DNA in *Xenopus* egg extracts supplemented either with an excess of recombinant 6xHis-tagged wild-type (WT) ubiquitin or various chain deficient mutants (Figure S3C). In the presence of WT ubiquitin, Dbn1 was heavily poly-ubiquitylated and migrated as different high molecular weight species on the gel (Figure S3C lane 1). In contrast, addition of a ubiquitin mutant unable to form lysine-linked chains (i.e. all lysines substituted with arginines, referred to as “noK”) resulted in faster migrating Dbn1 species consistent with conjugation of shorter ubiquitin chains or multiple mono-ubiquitins (Figure S3C lane 6). The high molecular weight species of Dbn1 were maintained in the presence of ubiquitin variants either unable to form K63-linked chains (“K63R”) or ubiquitin only capable of forming K48-linked chains (“K48only”) (Figure S3C lanes 3-4). Conversely, upon addition of a ubiquitin variant unable to form K48-linked chains (“K48R”) or ubiquitin able to form K63-linked chains only (“K63only”), the ubiquitylated Dbn1 species migrated faster, similar to that observed with the noK ubiquitin mutant (Figure S3C lanes 2, 5 and 6). Collectively, these data support that DSBs mainly induce K48-linked poly-ubiquitylation of Dbn1.

As K48-linked poly-ubiquitylation is a canonical signal for proteasomal degradation of the targeted protein (Chau et al., 1989; Swatek and Komander, 2016; Yau and Rape, 2016), we next examined whether DSB-induced ubiquitylation of Dbn1 would target the protein for degradation. Indeed, addition of proteasome inhibitor to egg extracts greatly stabilized ubiquitylated Dbn1 in the presence of DSB plasmid DNA (Figure S3D). Overall, we conclude that DSB-induced activation of the ATM/ATR-mediated DDR elicits K48-linked poly-ubiquitylation of the actin-organizing protein Dbn1, resulting in its proteasomal degradation.

### DSB-induced ubiquitylation of Dbn1 is mediated by the SCF^β-Trcp1^ ubiquitin E3 ligase

To elucidate the mechanism of DSB-induced Dbn1 ubiquitylation, we aimed at identifying the ubiquitin E3 ligase responsible for the modification. The largest family of ubiquitin E3 ligases are the Cullin-RING ligases (CRLs), which concurrently are known to primarily induce K48-linked poly-ubiquitylation of substrates resulting in their proteasomal degradation (Harper and Schulman, 2021; Petroski and Deshaies, 2005). We therefore reasoned that DSB-induced ubiquitylation of Dbn1 could be mediated by a CRL complex. To test this, we supplemented egg extracts with a pan-Cullin inhibitor (“Culi”) prior to induction of the DDR by DSB plasmid DNA and found that the inhibitor abolished DSB-induced ubiquitylation and stabilized the Dbn1 protein (Figure 3C).

In our UBIMAX analyses, we noted that ubiquitylation of Dbn1 followed a similar induction as ubiquitylation of Ku80 in response to DSBs (Figures 1I, 2D and 2I). Ku80 is known to be ubiquitylated by the Skp1-Cul1-Fbxl12 (SCF^Fbxl12^) complex in response to DSBs, triggering the dissociation of the Ku-complex from DNA (Postow and Funabiki, 2013; Postow et al., 2008). Consequently, we wondered whether the two proteins also shared the Cullin-dependent mechanism of ubiquitylation. To explore this, we supplemented egg extracts with recombinant dominant negative Cul1, Cul3, Cul4a and/or-b or Cul5 protein prior to stimulation with DSB plasmid DNA (Figure 3D). As expected, only dominant negative Cul1 abolished Ku80 ubiquitylation and stabilized the unmodified protein (Figure 3D lanes 4-6). In contrast, Cdt1, which is a known target of Cul4a^CDT2^ (Higa et al., 2006; Higa et al., 2003; Hu et al., 2004; Jin et al., 2006; Senga et al., 2006), was stabilized only in the presence of dominant negative Cul4a (Figure 3D lanes 10-12 and 16-18). Similar to Ku80, we observed loss of Dbn1 ubiquitylation and corresponding stabilization of the unmodified Dbn1 protein in the presence of dominant negative Cul1 (Figure 3D lanes 4-6). This was further validated by immunodepletion of Cul1 from egg extracts, which abolished ubiquitylation and completely stabilized both Ku80 and Dbn1 in the presence of DSB plasmid DNA (Figure S3E compare lanes 1-4 with 5-8).

Having established that Dbn1 is targeted for ubiquitylation in a Cul1-dependent manner we next sought to identify the Cul1 substrate targeting protein responsible for Dbn1 ubiquitylation. In addition to Cul1, Cul1 E3 ligase complexes consist of the RING-containing protein Rbx1, adapter protein Skp1 and a substrate targeting F-box protein (Feldman et al., 1997; Harper and Schulman, 2021; Skowyra et al., 1997). Since Dbn1 and Ku80 share the SCF-mediated mechanism of ubiquitylation upon DSBs, we first tested whether Dbn1 ubiquitylation also depended on the F-box protein Fbxl12. However, immunodepletion of Fbxl12 in egg extracts only caused stabilization of unmodified Ku80 but had no effect on DSB-induced ubiquitylation of Dbn1 (Figure S3E lanes 10-12). This suggests that the SCF complex utilizes different F-box proteins to recognize and target Ku80 and Dbn1 for DSB-induced ubiquitylation, respectively.

Considering that the SCF complex can interact with more than 70 F-box proteins (Yumimoto et al., 2020), we took advantage of a mass spectrometry-based approach to explore the mechanism for recognition and ubiquitylation of Dbn1 in response to DSBs (Figure 3E). As we found that DSB-induced ubiquitylation of Dbn1 required ATM activity (Figure 3B), we reasoned that a DSB-induced interaction between Dbn1 and the SCF complex would depend on ATM activity. Therefore, we performed a Dbn1 immunoprecipitation-mass spectrometry (IP-MS) experiment in which egg extracts were left untreated or supplemented with either ATM-or proteasome inhibitor before initiating a response by addition of either undamaged or DSB plasmid DNA (Figure 3E). Both *Xenopus laevis* isoforms of Dbn1 (Dbn1.S and Dbn1.L) were strongly enriched in quadruplicate Dbn1-immunoprecipitated samples compared to mock immunoprecipitation (Figure S3F). Consistent with DSB-induced ubiquitylation targeting Dbn1 for proteasomal degradation (Figure S3D), we detected an enriched interaction between Dbn1 and 14 proteasomal subunits in the presence of DSB and proteasome inhibitor from this unbiased proteomics approach (Figure 3F). Next, we examined the DSB-induced and ATM-dependent Dbn1 interactors, which revealed an enrichment of ubiquitin, Skp1, Cul1, Nedd8 and a single F-box protein, β-Trcp1 (Figure 3G). These proteins were specifically enriched in the presence of DSBs as compared to undamaged DNA, and each was significantly lost upon ATM inhibition (Figure 3H). Importantly, as the activity of Cullin ubiquitin E3 ligases requires neddylation of the Cullin subunit (Hori et al., 1999; Kamura et al., 1999; Read et al., 2000), the observed enrichment of Skp1-Cul1-β-Trcp1 along with ubiquitin and Nedd8 (Figure 3G) supports that Dbn1 interacts with the active SCF^β-Trcp1^ complex upon DSBs.

As our Dbn1 IP-MS experiment corroborated our previous finding that Cul1 is required for DSB-induced ubiquitylation of Dbn1 (Figure 3C-D), and further suggested that Dbn1 is ubiquitylated by the SCF^β-Trcp1^ complex upon DSBs (Figure 3E and 3G), we next investigated the requirement of the SCF substrate recognition factor β-Trcp1 for DSB-induced ubiquitylation of Dbn1. To this end, we raised two antibodies against *Xenopus laevis* β-Trcp1. However, as immunodepletion of β-Trcp1 could not be verified by WB using these antibodies, we instead confirmed their ability to recognize β-Trcp1 as well as to enrich the Skp1 and Cul1 components of the SCF complex from egg extracts by IP-MS (Figure S3G). Immunodepletion of β-Trcp1 using either antibody dramatically reduced DSB-induced ubiquitylation of Dbn1 (Figure S3H). We substantiated this by performing denaturing His-ubiquitin pulldowns after addition of recombinant 6xHis-tagged ubiquitin and DSB plasmid DNA to egg extracts, which showed a complete loss of Dbn1 ubiquitylation upon either Cul1 or β-Trcp1 immunodepletion (Figure S3I). Critically, DSB-induced Dbn1 ubiquitylation was restored by addition of recombinant β-Trcp1 protein to β-Trcp1 immunodepleted egg extracts (Figures 3I and S3J), demonstrating the specific requirement for β-Trcp1 for DSB-induced ubiquitylation of Dbn1. Dbn1 ubiquitylation was also observed in HeLa cells 30 minutes after DSB formation induced by treatment with ionizing radiation (IR) (Figure 3J). The IR induced DBN1 signal observed was highly reduced upon siRNA-mediated knock-down of DBN1 in HeLa cells (Figures S3K), confirming that DNA damage-induced ubiquitylation of DBN1 also occurs in human cells. As observed in egg extracts, DNA damage-induced DBN1 ubiquitylation was fully dependent on ATM- and Cullin ubiquitin E3 ligase activity (Figure 3K). In addition, knock-down of β-Trcp1 by two independent siRNAs also eliminated DBN1 ubiquitylation in response to IR (Figure 3L). In summary, these data establish a conserved mechanism in which the SCF^β-Trcp1^ complex mediates the DSB-induced and ATM-dependent ubiquitylation of Dbn1 identified by UBIMAX.

### DSB-induced Dbn1 ubiquitylation is driven by a DDR-specific β-Trcp1 degron

To gain further mechanistic insight to the DDR-dependent and SCF^β-Trcp1^-mediated ubiquitylation of Dbn1, we sought to identify a putative β-Trcp1 degron in the Dbn1 protein sequence. β-Trcp1 recognizes its substrates via a [D/E/S]-[S/D/E]-G-X-X-[S/E/D] degron motif, in which phosphorylation of both the flanking serine residues is required (Frescas and Pagano, 2008; Margottin et al., 1998). Indeed, upon scanning the *Xenopus laevis* Dbn1 sequence, we found a putative motif, S-E-G-Y-F-S (amino acids 604-609) located in the unstructured C-terminal region of Dbn1, which is fully conserved across different vertebrate species (Figure 4A). Intriguingly, the last serine residue in this putative β-Trcp1 degron also forms part of a double ATM consensus [S/T]-Q phosphorylation motif (S609 and S611, respectively) (Kim 1999, O’Neill 2000). Our Dbn1 IP-MS experiment described above (Figure 3E) corroborate the assumption that these residues are phosphorylated in a DNA damage and ATM-dependent manner, as we abundantly detected an unmodified Dbn1 peptide containing this putative β-Trcp1 degron in the undamaged condition, while addition of DSB plasmid DNA to egg extracts abrogated detection of this peptide (Figure S4A, right). Moreover, supplementing egg extracts with an ATM inhibitor prior to the DSB stimulus reenabled the detection of the unmodified peptide. The lack of detection was not due to the general Dbn1 sequence context, as the upstream peptide was detected equally across all conditions (Figure S4A, left). Although the phosphorylated peptide was not detected by MS, we speculate that DSB-induced and ATM-mediated phosphorylation of these SQ motifs could be occurring, and concomitantly enable recognition of Dbn1 by β-Trcp1 specifically in response to DDR activation To further investigate the phosphorylation status of the SQ motifs situated in direct connection with the putative β-Trcp1 degron in the Dbn1 C-terminus, we raised a phospho-specific antibody against these serine residues (Dbn1-pS609/pS611). Using this antibody, we confirmed phosphorylation of these Dbn1 SQ motifs as early as 5 min following stimulation with DSB plasmid DNA (Figure 4B, lanes 1-6). While inhibition of either ATM, ubiquitin E1 enzyme or Cullin E3 ligases prevented ubiquitylation of Dbn1, Dbn1-S609/S611 phosphorylation was heavily reduced upon ATM inhibition but remained unaffected by inhibition of ubiquitin E1 enzyme or Cullin E3 ligases (Figure 4B). Collectively, this suggests that Dbn1-S609/S611 phosphorylation is mediated by the apical DDR kinase ATM and occurs upstream of Dbn1 ubiquitylation.

**Figure 4.**
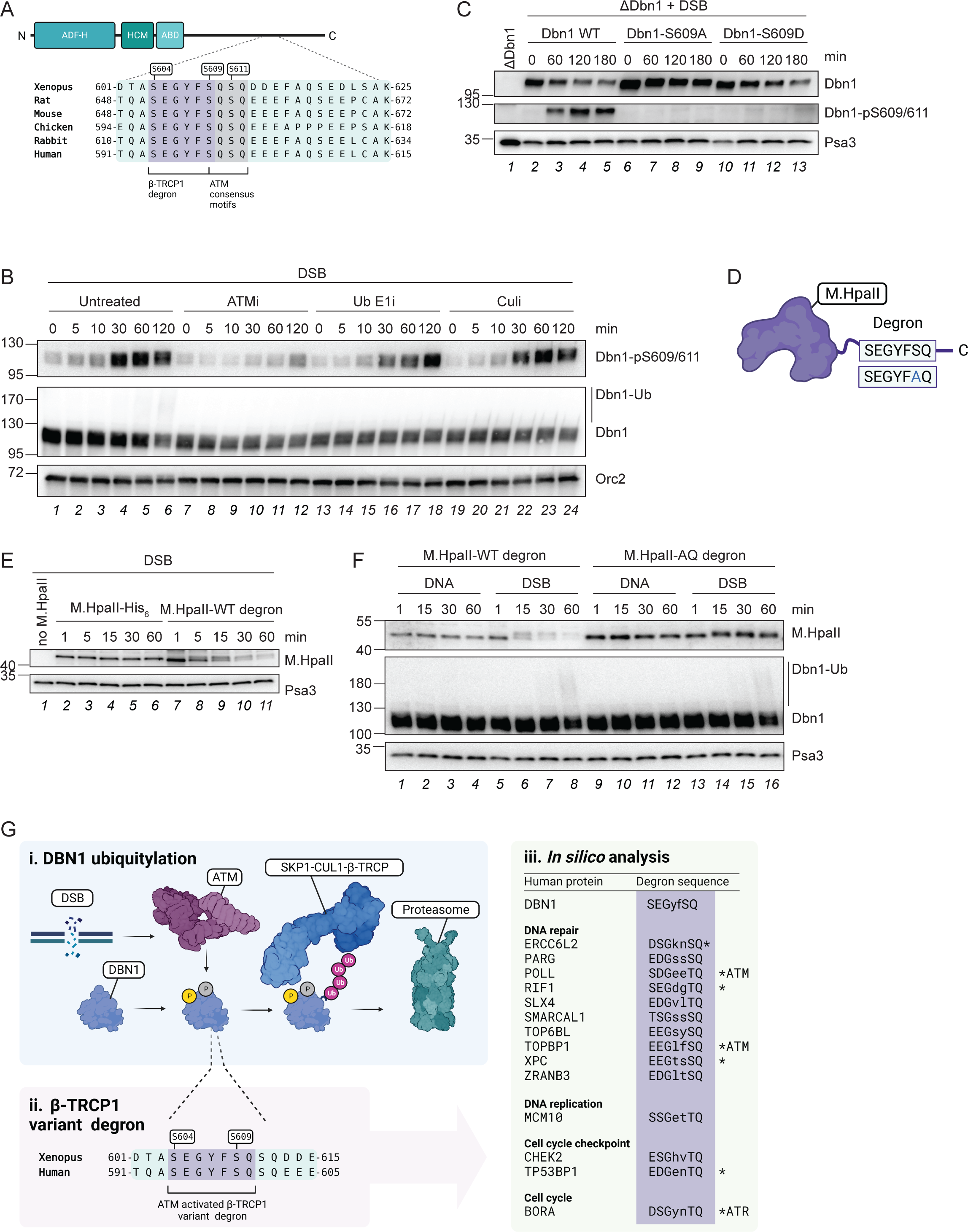
A variant β-Trcp1 degron is necessary and sufficient for inducing Dbn1 and general protein degradation in response to DSBs. **A.** Schematic representation of the variant β-Trcp1 degron located in the Dbn1 C-terminus and conserved across vertebrate species. **B.** Egg extracts were left untreated or supplemented with ATM inhibitor (“ATMi”), ubiquitin E1 inhibitor (“Ub E1i”), or neddylation E1 inhibitor (“Culi”), prior to addition of linearized plasmid DNA (“DSB”). Samples were transferred to sample buffer at the indicated timepoints and analysed by WB using antibodies against Dbn1 phosphorylated at serine residues 609/611 (Dbn1-pS609/S611) and total Dbn1. Orc2 served as a loading control. **C.** Recombinant Dbn1 WT, S609A, or S609D, were mixed with Dbn1-immunodepleted egg extracts. Samples were collected from Dbn1-immunodepleted egg extract prior to addition of recombinant protein, and at the indicated timepoints following addition of protein and linearized plasmid DNA. Samples were analysed by WB using antibodies against Dbn1 and Dbn1-pS609/611. Psa3 served as a loading control. **D.** Schematic representation of the recombinant proteins generated by insertion of the WT and mutated variant β-Trcp1 degron at the M.HpaII C-terminus. **E.** Recombinant M.HpaII protein with or without the variant β-Trcp1 degron or buffer (“no M.HpaII”) was mixed with egg extracts prior to addition of linearized plasmid DNA. Samples were transferred to sample buffer at the indicated timepoints and analysed by WB using antibodies against M.HpaII. Psa3 served as a loading control. **F.** Recombinant M.HpaII protein with the WT or AQ-mutated variant β-Trcp1 degron was mixed with egg extracts prior to addition of undamaged-(“DNA”) or linearized plasmid DNA (“DSB”). Samples were transferred to sample buffer at the indicated timepoints and analysed by WB using antibodies against M.HpaII and Dbn1. Psa3 served as a loading control. **G.** i) DNA damage, such as DSBs, activates the apical DDR kinase ATM, which mediates the phosphorylation of the actin-organizing protein Dbn1 at the S609 SQ site. This primes the conserved variant degron with which it is connected (ii) for recognition by the F-box protein β-Trcp1, resulting in ubiquitylation by the SCF^β-Trcp1^ E3 ligase compelx and subsequent proteasomal degradation of the Dbn1 protein. iii) *In silico* analysis of the human proteome reveals numerous proteins containing a potential ATM/ATR-activated β-Trcp1 degron, a subset of which are involved in the DNA damage response. * indicates proteins for which the S/TQ site of the putative variant β-Trcp1 degron is known to be phosphorylated.

To investigate whether phosphorylation of the SQ motif situated in the putative β-Trcp1 degron is required for Dbn1 ubiquitylation, we produced *in vitro* translated recombinant WT, phosphodeficient (S609A) and phosphomimic (S609D) Dbn1 proteins from rabbit reticulocyte lysates (Figure 4A). We reasoned that if S609-phosphorylation was required for Dbn1 ubiquitylation, the S609A mutation should render Dbn1 refractory to ATM-mediated phosphorylation and thus preclude recognition by β-Trcp1 for SCF^β-Trcp1^-mediated ubiquitylation in response to DSBs. In contrast, the Dbn1-S609D phosphomimic mutant would not require DSB-induced and ATM-dependent phosphorylation for SCF^β-Trcp1^-mediated ubiquitylation. To test this hypothesis, we first confirmed that the recombinant WT Dbn1 protein was subjected to ATM-dependent phosphorylation in response to DSBs in egg extracts immunodepleted for endogenous Dbn1 (Figure S4B). However, while recombinant Dbn1 WT and -S609A proteins were readily produced by *in vitro* translation in rabbit reticulocyte lysates, we were initially not able to produce the Dbn1-S609D mutant. Interestingly, addition of Cullin ubiquitin E3 ligase inhibitor to the *in vitro* translation reaction enabled production of Dbn1-S609D protein, while addition of proteasome inhibitor resulted in a heavily modified form of Dbn1-S609D (Figure S4C). These observations suggest that recombinant Dbn1-S609D is spontaneously ubiquitylated by a Cullin E3 ligase and subsequently degraded by the proteasome in rabbit reticulocyte lysates, thus hinting at a conserved mechanism for Dbn1 ubiquitylation. To test the relevance of these Dbn1 variants in egg extracts, we supplemented Dbn1-immunodepleted extracts with either the WT, phosphodeficient or phosphomimic recombinant Dbn1 proteins. Upon DSB addition, WT Dbn1 was phosphorylated at the double SQ motifs and the amount of unmodified protein correspondingly declined over time (Figure 4C lanes 2-5). This correlated with WT Dbn1 ubiquitylation upon addition of the DSB plasmid (Figure S4D, lanes 3-4). Importantly, the Dbn1-S609A phosphodeficient mutant was noticeably stabilized in the presence of DSBs (Figure 4C lanes 6-9) and no damage-induced ubiquitylation was observed (Figure S4D, lanes 7-8). In contrast, the Dbn1-S609D phosphomimic mutant was unstable despite lacking S609/S611 phosphorylation (Figure 4C lanes 10-13). We also confirmed that this phosphorylation event is conserved and stimulated by DNA damage in HeLa cells transiently transfected with either GFP-tagged WT DBN1 or the corresponding phosphodeficient GFP-DBN1-S599A mutant (Figure S4E).

As recognition of target proteins by β-Trcp1 requires phosphorylation of both serine residues of the target protein degron (Dbn1-S604 and -S609) (Frescas and Pagano, 2008; Margottin et al., 1998), we produced the corresponding single and double phosphodeficient Dbn1 mutants as well as mutants of the second SQ motif immediately downstream of the putative β-Trcp1 degron (S611) (Figure S4F). Mutation of either S604 or S609 rendered recombinant Dbn1 completely stable, despite the S604 mutation was permissive to DSB-induced S609/S611 phosphorylation (Figure S4F lanes 1-4 and 13-16). Nor did S611A mutation affect the complete stabilization of Dbn1 conferred by the S609A mutation (Figure S4F lanes 5-12). Together, these data demonstrate that Dbn1 is targeted for ubiquitylation by the SCF^β-Trcp1^ ligase complex through recognition of a variant β-Trcp1 degron, SEGYFSQ, which is specifically sensitive to DNA damage through the direct incorporation of an ATM consensus phosphorylation site.

### The Dbn1 degron represents a DDR-sensitive β-Trcp1 variant degron

We wondered whether the DDR-sensitive β-Trcp1 variant degron identified in the Dbn1 C-terminus could function as a general motif for conveying DNA damage-induced ubiquitylation and degradation via the SCF^β-Trcp1^ ubiquitin ligase. To investigate this, we cloned the Dbn1 S-E-G-Y-F-S-Q motif onto the C-terminus of the *Haemophilus parainfluenzae* methyltransferase M.HpaII, a protein not native to *Xenopus* egg extracts (Figure 4D). Strikingly, while the recombinant WT M.HpaII protein was stable in egg extracts challenged by DSB plasmid DNA, the M.HpaII protein tagged with the Dbn1 degron exhibited a mass shift as well as a gradual disappearance of the protein, suggesting the occurrence of DSB-induced phosphorylation and subsequent degradation (Figure 4E). To confirm degron-targeted degradation of M.HpaII upon DSBs, we additionally cloned the phosphodeficient mutant motif, S-E-G-Y-F-A-Q onto the M.HpaII C-terminus (Figure 4D) and analysed the stability of the degron-tagged M.HpaII proteins in the presence of either undamaged or DSB plasmid DNA in egg extracts (Figure 4F). From this, we observed that M.HpaII tagged with the WT Dbn1 degron was destabilized in response to DSB plasmid DNA but remained stable in the presence of undamaged plasmid (Figure 4F lanes 1-8), confirming that the Dbn1 degron conferred DSB-specific targeting of the M.HpaII protein. Remarkably, M.HpaII tagged with the phosphodeficient Dbn1 degron remained unmodified and stable both in the presence of undamaged and DSB plasmid DNA (Figure 4F lanes 9-16).

To test whether introduction of the Dbn1 degron onto the C-terminus of M.HpaII confers degradation by the same mechanism as Dbn1, we monitored M.HpaII protein modification and stability upon DSBs in the presence or absence of ATM-or Cullin E3 ligase inhibitors (Figure S4G). Indeed, ATM inhibition abolished phosphorylation of degron-tagged M.HpaII and stabilized the protein in the presence of DSBs, whereas Cullin E3 ligase inhibition was permissive to M.HpaII phosphorylation and stabilized the phosphorylated protein. Collectively, this confirms that the DDR-sensitive β-Trcp1 degron identified in Dbn1 is transferrable and sufficient for conferring ATM-dependent phosphorylation and subsequent recognition by the SCF^β-Trcp1^ ubiquitin ligase complex.

## Discussion

As ubiquitylation is a key signaling modulator involved in regulating most cellular functions, methods for global, unbiased profiling of ubiquitylation events are needed. Here, we presented a new method, UBIMAX, which efficiently and specifically identifies dynamic and quantitative protein ubiquitylation under defined and adaptable conditions of choice in *Xenopus* egg extracts. We demonstrate that UBIMAX can detect highly DNA-damage specific ubiquitylation events and identify the previously uncharacterized, DSB-induced ubiquitylation of the actin-organizing protein, Dbn1. We unravel the conserved mechanism for this ubiquitylation event and show that it is mediated by the SCF^β-Trcp1^ E3 ligase and depends on direct ATM-dependent phosphorylation of a variant β-Trcp1 degron (Figure 4G). We further show that this variant β-Trcp1 degron is necessary and sufficient for DSB-induced degradation of a model substrate, M.HpaII. Collectively, our work demonstrates UBIMAX’s capacity to identify novel and conserved mechanisms of the ubiquitylation response to a defined DNA lesion.

In this study, we have used UBIMAX to investigate ubiquitylation dynamics in response to DNA damage and identify protein ubiquitylation specifically induced by DSBs or DPCs. By detecting proteins previously known to be ubiquitylated upon DNA damage (Figure 1I) and validating the previously unknown ubiquitylation of the actin-organizing protein Dbn1 (Figures 2J and 3A), we show that the denaturing ubiquitin enrichment approach we have utilized for UBIMAX is successful in specifically enriching for ubiquitin-conjugated proteins while eliminating ubiquitin-interacting proteins. This is due to the ease of supplementing *Xenopus* egg extracts with, in this case, recombinant His6-tagged ubiquitin protein. Another advantage of the *Xenopus* egg extract system is the possibility to generate site-specific DNA lesions of interest and follow the response to these lesions with temporal precision. We have shown that UBIMAX is capable of detecting ubiquitylation events specific to such DNA lesions as well as identifying common DNA damage-related ubiquitylation responses (Figures 1I, 2B-C and S2B). Moreover, due to the synchronous nature of *Xenopus* egg extracts and the ability to easily inhibit or immunodeplete essential proteins, UBIMAX could be utilized to interrogate the ubiquitylation response to defined processes such as DNA replication and mitosis with temporal precision and in the absence of essential protein functions. Furthermore, we found evidence to suggest that UBIMAX can be utilized to investigate other aspects of ubiquitin signaling that have remained technically challenging. For example, we detected DNA damage-induced ubiquitylation of the ubiquitin E3 ligases Chfr and Hltf (Figures 1I and 2B-C), which auto-ubiquitylate in response to DNA damage (Chaturvedi et al., 2002; Lin et al., 2011; Liu et al., 2013). In addition to Chfr and Hltf, we observed ubiquitylation of a further 12 ubiquitin E3 ligases in response to DSBs, DPCs or both (Tables S1 and S3). This indicates the potential of UBIMAX to profile active ubiquitin ligases in response to a condition of choice by detecting ubiquitin E3 ligase auto-ubiquitylation. By further taking advantage of the possibility to immunodeplete or chemically inactivate specific ubiquitin E3 ligases in egg extract, UBIMAX could be utilized for interrogating ligase-substrate relationships. Finally, we envision that UBIMAX could be employed to provide linkage-specific information about global ubiquitylation as well as be adapted to investigate other ubiquitin-like protein modifications via the direct addition of recombinant 6xHis-tagged linkage-specific ubiquitin mutants or ubiquitin-like proteins to *Xenopus* egg extracts.

Using UBIMAX, we detected the novel DSB-induced ubiquitylation of the actin-organizing protein Dbn1 (Figure 1I). Dbn1 binds to and stabilizes actin filaments by preventing depolymerization of actin subunits from the filament barbed end, inhibiting Cofilin-mediated filament severing and inducing actin filament bundling (Grintsevich and Reisler, 2014; Mikati et al., 2013; Sharma et al., 2012; Sharma et al., 2011; Worth et al., 2013). Actin-binding proteins have been shown to associate with DSBs and actin filament polymerization was proposed to regulate DSB localization and repair by homologous recombination (HR) in a manner depending on the DDR (Aymard et al., 2017; Belin et al., 2015; Caridi et al., 2018; Schrank et al., 2018; Zagelbaum et al., 2023). However, how the DDR connects to and regulates actin filament dynamics has remained elusive and Dbn1 has not previously been shown to be involved in DSB repair. We show that the apical DDR kinase, ATM, by inducing phosphorylation of Dbn1 in the presence of DSBs, triggers the ubiquitylation and degradation of the Dbn1 protein (Figures 4B). Furthermore, our Dbn1 IP-MS data indicate that the interaction between Dbn1 and actin filament-related factors (e.g. capza1, capzb) depend on ATM activity while the interaction between Dbn1 and actin (acta1) is reduced in the presence of DSBs (Figure 3G). It would be interesting to understand if DDR-induced degradation of Dbn1 directly impacts actin filament dynamics and thereby DSB repair.

We show that Dbn1 is ubiquitylated by the SCF^β-Trcp1^ E3 ligase through recognition of a variant β-Trcp1 degron (S-E-G-Y-F-S-Q) situated in the unstructured Dbn1 C-terminus (Figures 3 and 4C). Recognition requires phosphorylation of both serine residues, of which the one situated in the SQ motif depends on ATM activity (Figures 4B-C and S4F). We further show that this variant β-Trcp1 degron is necessary and sufficient for degradation of a model substrate, M.HpaII, upon addition of DSBs (Figures 4E-F). We note, however, that DSB-induced degradation of the model substrate occurred with faster kinetics than Dbn1 (compare Figures 4B and E). This indicates that additional regulatory mechanisms exist for Dbn1 ubiquitylation. Indeed, the Dbn1 sequence contains additional conserved S/TQ motifs upstream and immediately downstream of the β-Trcp1 degron. Together, these potential ATM phosphorylation sites could form an S/TQ cluster domain, for which it has been suggested that all S/TQ sites need to be phosphorylated in order for the domain to adopt a structure permissive for DNA damage-induced protein-protein interactions (Traven and Heierhorst, 2005). On the other hand, this S/TQ cluster could stimulate phosphatase recruitment and thereby counteract the activation of the degron (Lee and Chowdhury, 2011). In fact, the immediate downstream SQ site (S611) has previously been described as phosphorylated in an ATM-dependent manner in response to oxidative stress in C.*elegans* (Dbn1-S647), stabilizing the protein, and dephosphorylated by the PTEN phosphatase (Kreis et al., 2019; Kreis et al., 2013). Finally, Dbn1 is suggested to exist in a closed confirmation, which is alleviated by cyclin-dependent kinase-like 5 (Cdk5)-mediated phosphorylation of S142 allowing access to the C-terminus (Worth et al., 2013). Indeed, in our Dbn1 IP-MS experiment we detected the phosphorylation of this site in *Xenopus* egg extracts (Table S4). Thus, we hypothesize that these additional conserved S/TQ sites surrounding the degron may act as additional regulatory elements to finetune Dbn1 protein stability and actin filament organization.

As the β-Trcp1 degron identified in Dbn1 induced DSB-dependent degradation of a M.HpaII model substrate (Figures 4E-F), we wondered if this variant β-Trcp1 degron could represent a general mechanism for inducing SCF-mediated ubiquitylation and degradation of proteins in response to DDR activation. We further envision that this variant β-Trcp1 degron could be used to induce specific and timely degradation of essential proteins in response to DSB addition to e.g. investigate specific DNA repair pathways. While a functional β-Trcp1 degron containing an ATR-regulated phosphorylation site has been reported for the mitotic regulator BORA in human cells (Qin et al., 2013; Seki et al., 2008), the general occurrence of such a variant degron has not been previously described. To assess the global distribution of such a variant DDR-β-Trcp1 degron, we carried out an *in silico* analysis by searching the human proteome for the occurrence of the motif, [D/E/S/T]-[D/E/S]-G-X-X-[S/T]-Q (Table S6). This analysis identified close to 300 proteins displaying DDR-β-Trcp1 variant degrons, of which we noted several involved in regulating DNA repair, checkpoint, replication, and cell cycle pathways (Figure 4Giii). Despite that only the identified degrons of BORA, TOPBP1 and POLL are known to be phosphorylated by the apical DDR kinases ATM or ATR (Qin et al., 2013; Sastre-Moreno et al., 2017; Yamane et al., 2002), our *in silico* analysis show that 41 proteins have previously been reported phosphorylated at the S/TQ motif of the putative DDR-β-Trcp1 degrons. While further studies are required to determine whether these putative degrons are functional, we find evidence for a wider existence of a DDR-responsive β-Trcp1 degron. We could envision that such a degron could provide a hitherto unappreciated mechanism for inducing a coordinated SCF^β-Trcp1^-mediated ubiquitylation program in response to DNA damage and thus regulate DSB repair, DDR-, and checkpoint activity.

## Materials and methods

### Xenopus egg extracts and reactions

Egg extracts were prepared using *Xenopus laevis* (Nasco Cat #LM0053MX, LM00715MX). All experiments involving animals were approved by the Danish Animal Experiments Inspectorate and are conform to relevant regulatory standards and European guidelines. Preparation of Xenopus high speed supernatant interphase egg extracts (HSS) was performed as described previously (Lebofsky et al., 2009). Reactions were performed at room temperature (RT) using HSS supplemented with 3 μg/mL nocadazole and ATP regeneration mix (20 mM phosphocreatine, 2 mM ATP, 5 μg/mL creatine phosphokinase). Where indicated, HSS was supplemented with various inhibitors and incubated for 10 or 20 minutes at RT prior to addition of plasmid DNA. To block de novo ubiquitylation, egg extracts were supplemented with 200 µM Ubiquitin E1 inhibitor (MLN7243, Active Biochem). Activity of the apical DNA damage response kinases were inhibited using ATM inhibitor (KU-559333, Selleckchem), ATR inhibitor (AZ20, Sigma-aldrich) and DNA-PKcs inhibitor (NU7441, Selleckchem) at final concentrations of 100 µM. Cullin E3 ligase activity was blocked by supplementing egg extracts with 100 µM neddylation E1 enzyme inhibitor (MLN4924, R&D systems). Proteasome activity was inhibited via addition of 200 μM MG262 (Boston Biochem). Egg extracts were supplemented with recombinant proteins as detailed below and incubated for 10 min at RT before addition of plasmid DNA unless otherwise stated. Where indicated, 6xHis-tagged human recombinant ubiquitin (Boston Biochem) was added to egg extracts at a final concentration of 0.1 μg/μL unless otherwise stated. To investigate ubiquitin-conjugation linkage type, 6xHis-tagged ubiquitin mutants (Boston Biochem) were added at final concentrations of 1 µg/µL. For testing the Cullin E3 ligase specificity of Dbn1 ubiquitylation, egg extracts were supplemented with recombinant dominant negative *Xenopus* Cul1, Cul3, Cul4a, Cul4b, and Cul5 proteins at final concentrations of 0.3 µg/µL, 0.2 µg/µL, 1.1 µg/µL, 0.3 µg/µL, and 0.3 µg/µL, respectively. Where indicated, *in vitro* translated *Xenopus* β-Trcp1, WT and mutant Dbn1 proteins were generally added to egg extracts in a 1:4-6.25 or 1:10 ratio for reticulocyte and wheat germ systems, respectively (see further details below). For testing ubiquitylation of a model substrate, recombinant *Haemophilus parainfluenzae* methyltransferase M.HpaII without or with the WT or AQ mutated β-Trcp1 degron identified in Dbn1 were added to egg extracts in a 1:10 ratio and incubated for 10 or 30 min at RT before addition of plasmid DNA. Reactions were initiated by addition of 15 ng/μL plasmid DNA substrate as indicated.

### Preparation of DNA substrates

The DSB-mimicking plasmid DNA substrate was generated by linearizing pBlueScript II KS (pBS) through enzymatic digestion using XhoI. Circular pBS was used as the undamaged control. To generate a radiolabeled DSB substrate, pBS was first nicked with nb.BsrDI and subsequently radiolabeled with [α-^32^P]dATP via nick translation synthesis by DNA Pol I for 20 min at 16°C. Radiolabeled pBS was subsequently linearized as described above.

ssDNA-DPC was previously described in (Larsen et al., 2019) as pDPC^ssDNA^. To generate SSB-DPC, we first created pFRT by inserting the specific Flp recognition target site sequence into pBS, by replacing the EcoRI-HindIII fragment with the sequence 5’-AAT TCG ATA AGT TCC TAT TCG GAA GTT CCT ATT CTC TAG AAA GTA TAG GAA CTT CAT CA-3’. For the crosslinking reaction, pFRT was mixed with Flp-nick-His6 in reaction buffer (50 mM Tris-HCl pH 7.5, 50 mM NaCl, 20 µg/ml BSA and 1 mM DTT) and incubated overnight at 30°C (Nielsen et al., 2009).

### Antibodies, Immunodepletion and -detection

Antibodies against *Xenopus* Mcm6 (Semlow et al., 2016), Orc2 (Fang and Newport, 1993), Cdt1 (Arias and Walter, 2005) as well as M.HpaII (Larsen et al., 2019) were previously described. The antibody against His (631212, Fisher Scientific) is commercially available. The following antibodies were raised against the indicated peptides derived from *Xenopus laevis* proteins (New England Peptide now Biosynth): Dbn1 (Ac-CWDSDPVMEEEEEEEEGGGFGESA-OH), Ku80 (CMEDEGDVDDLLDMM), Cul1 (H2N-MSSNRSQNPHGLKQIGLDQC-amide), Fbxl12 (Ac-CRGIDELKKSLPNSKVTN-OH), Psa3 (Ac-CKYAKESLEEEDDSDDDNM-OH), β-Trcp1-INT (Ac-GQYLFKNKPPDGKTPPNSC-amide), β-Trcp1-NT (H2N-MEGFSSSLQPPTASEREDC-amide), and Dbn1-pS609/611 (Ac-CSEGYF(pS)Q(pS)QDED-amide).

Antibodies against human proteins used in this study include ubiquitin (P4D1 (sc-8017), Santa Cruz), CHK1-pS345 (#2341, Cell Signaling), DBN1 (TA812128, Thermo Fisher Scientific), CUL4A (2699S, Cell Signaling), GAPDH (sc-20357 HRP, Santa Cruz), all of which are commercially available.

To immunodeplete *Xenopus* egg extracts, Protein A Sepharose Fast Flow (PAS) (GE Health Care) beads were bound to the indicated antibodies with a stock concentration of 1 mg/mL and at a beads-antibody ratio of 1:4 overnight at 4°C. Beads were then washed twice with 500 μl PBS, once with ELB buffer (10 mM HEPES, pH 7.7; 50 mM KCl; 2.5 mM MgCl2; and 250 mM sucrose), twice with ELB buffer supplemented with 0.5 M NaCl, and twice with ELB buffer. One volume of HSS was then depleted by addition of 0.2 volumes of antibody-bound beads and incubating at RT for 15 minutes with end-over-end rotation, before being harvested. This was repeated one additional round for depletion of Dbn1 and two additional rounds for depletion of Cul1, Fbxl12, and β-Trcp1. Unless otherwise stated, the β-Trcp1-NT antibody was used for depletion of β-Trcp1.

For WB analysis, samples were added to 2x Laemmli sample buffer and resolved on SDS-PAGE gels. Proteins were visualized by incubation with the indicated antibodies and developed using the chemiluminescence function on an Amersham Imager 600 (GE Healthcare) or a Compact 2 developer (Protec). Specifically for WB analysis of *Xenopus* Dbn1 and human DBN1, the commercially available DBN1 antibody was used in Figures 2J, 3C-D, 3J-L, S3E, S3K, and S4E, while the antibody raised against *Xenopus* Dbn1 was used in Figures 3A-B, 3I, 4B-C, 4F, S3A-D, S3H-J, S4B-D, and S4F-G.

### Protein expression and purification

*Xenopus laevis* Dbn1 (L. homolog, Thermo) and β-Trcp1 (S. homolog encoded by bPZ934, a kind gift from Philip Zegerman (Collart et al., 2017)) was cloned into the pCMV-Sport vector under the Sp6 promoter. The Dbn1 mutant sequences were generated using the KOD Hot Start DNA Polymerase kit (Sigma-Aldrich), according to the manufacturer’s instructions. The proteins were then expressed by in vitro translation in rabbit reticulocytes lysate. Specifically, two reactions containing 40 μL TnT SP6 Quick Master Mix (Promega), 2 μL of 1 mM methionine and 1 μg of pCMV-Sport were incubated for 90 minutes at RT. For expression of Dbn1-S609D, this reaction was further supplemented with 200 µM neddylation E1 enzyme inhibitor. As a negative control for rescue experiments, a reaction without DNA was performed. The two reactions were subsequently mixed and concentrated at 4°C through an Amicon Ultra-0.5 Centrifugal Filter Unit (Millipore) with a 30 kDA cutoff to a total volume of 50 μL. The recombinant proteins used in the experiments presented in Figures 3I, 4C, S3J, S4D, and S4F were produced in this manner and added to egg extract in ratios of 1:4, 1:5, 1:4, 1:4, 1:6.25, respectively. Additionally, for the expression of Dbn1 WT and mutant proteins used in the experiment presented in Figure S4B, the TnT SP6 High-Yield Wheat Germ Protein Expression System (Promega) was used. The proteins were expressed according to the manufacturer’s instructions and added to egg extracts at a 1:10 ratio.

Plasmids expressing the dominant negative Cullin proteins, Cul1-NT, Cul3a-NT and Cul5a-NT C-terminally tagged with His_6_-and FLAG tags, were kind gifts from Prof. Alex Bullock and were expressed and purified as previously described (Canning et al., 2013). *Xenopus* gene fragment coding for Cul4b-NT (amino acids 159-510) was synthesised based on Xenbase sequences with addition of His_6_-and FLAG tag at C-termini, and cloned into pET28 and pET23 vectors, respectively. Cul4a-NT (amino acids 1-396) coding sequence was amplified from *Xenopus* cDNA and cloned into pET23 vector. Both proteins were purified as above.

M.HpaII-His_6_ was expressed and purified as previously described (Duxin et al., 2014). To generate the M.HpaII-WT degron and -AQ degron proteins, M.HpaII was first cloned into pHis_6_-SUMO (Liu et al., 2021) using primers 5’-TATAGGATCCATGAAAGATGTGTTAGATGATA-3’ and 5’-TATAGAGCTCTCAttcatgccattcaatcttctg-3’, the latter of which contains the sequence for the AviTag. Plasmid encoding M.HpaII-WT degron and - AQdegron were then constructed by addition of the Dbn1 WT or S609A degron sequence, S-E-G-Y-F-S/A-Q, to the C-terminus of pHis_6_-SUMO-M.HpaII-Avitag via PCR using primers 5’-atgcGCTAGCGGATCGGACTCA-3’ and 5’-tataGGTACCTTGGCTGAAATATCCTTCACTggattggaagtacaggttctcaa-3’ or 5’-tataGGTACCTTGGGCGAAATATCCTTCACTggattggaagtacaggttctcaa-3’ for WT and AQ mutant, respectively. The degron-tagged M.HpaII protein was subsequently expressed and purified as described in (Duxin et al., 2014) and the N-terminal His_6_-SUMO-tag cleaved off using the SUMO protease Ulp1.

### DNA repair assay

For assaying DSB repair in *Xenopus* egg extract, HSS was supplemented with the indicated inhibitors and proteins and reactions initiated by addition of radiolabelled linearized plasmid DNA. At the indicated timepoints, 2 µL reaction was added to 10 volumes of transparent stop buffer (50 mM TrisHCl, pH 7.5, 0.5% SDS, 25 mM EDTA), treated with 1 µL RNase A (Thermo) for 30 minutes followed by 1 µL Proteinase K (20 mg/mL, Roche) for 1 hour at 37°C. The DNA was purified by phenol/chloroform extraction, ethanol precipitated in the presence of glycogen (20 mg/mL, Roche), and resuspended in 10 μLubi of 10 mM Tris buffer (pH 7.5). The DNA was separated by 0.9% native agarose gel electrophoresis and visualized using a phosphorimager. Radioactive signal was quantified using ImageJ (NIH, USA) and quantifications graphed using Prism (GraphPad Software).

### Denaturing His-ubiquitin pulldown

To enrich ubiquitin-conjugated proteins, Ni-NTA superflow agarose beads (Qiagen) were washed thrice in denaturing pulldown buffer (6M Guanidine hydrochloride, 0.14 M NaH_2_PO_4_, 4.2 mM Na_2_HPO_4_, 10 mM Tris pH 7.8). At the indicated timepoints, *Xenopus* egg extract reactions supplemented with a final concentration of 0.1 µg/µL His_6_-ubiquitin were added to 4 volumes of beads for UBIMAX and 3.3 volumes for WB analysis, respectively, in a total of 50 volumes of denaturing pulldown buffer supplemented with 25 mM imidazole and 6.25 mM β-mercaptoethanol and incubated for 60 minutes or overnight at 4°C with end-over-end rotation. Beads were washed thrice in denaturing pulldown buffer supplemented with 10 mM imidazole and 5 mM β-mercaptoethanol, five times in wash buffer 2 (8 M urea, 78.4 mM NaH_2_PO_4_, 21.6 mM Na_2_HPO_4_, 10 mM Tris, pH 6.3) and twice in wash buffer 3 (8 M urea, 6.8 mM NaH_2_PO_4_, 93.2 mM Na_2_HPO_4_, 10 mM Tris, pH 8). For elution of ubiquitin-conjugated proteins for WB analysis, beads were resuspended in 2x Laemmli sample buffer with 0.5 M EDTA, boiled at 95°C for 5 min, and eluates separated from the beads by centrifugation through homemade nitex columns.

#### UBIMAX

For MS-based analysis of ubiquitin-conjugated proteins via UBIMAX, ubiquitylated proteins were enriched by denaturing His-ubiquitin pull down as described above. Beads were resuspended in 150µL wash buffer 3 and diluted with two volumes of 10 mM Tris pH 8.5 prior to on-bead digestion of proteins via addition of 500 ng modified sequencing grade Trypsin (Sigma) with incubations of 1 hour at 4°C and then overnight at RT with continuous mixing at 1200 rpm. Eluates containing digested peptides were separated from beads by centrifugation through a 0.45 µM PVDF filter column (Millipore) and cysteines were subsequently reduced and alkylated by addition of 5 mM TCEP and 10 mM CAA for 30 min at 30°C. Tryptic peptides were acidified with 10% trifluoroacetic acid (pH < 4) and purified by C18 StageTips prepared in-house (Rappsilber et al., 2003). Four plugs of C18 material (Sigma-Aldrich, Empore™ SPE Disks, C18, 47 mm) were layered per StageTip and activated in 100% methanol, then equilibrated in 80% acetonitrile 10% formic acid, and finally washed twice in 0.1% formic acid. Acidified samples were loaded on the equilibrated StageTips and washed twice with 50 mL 0.1% formic acid. StageTips were eluted into LoBind tubes with 80 mL of 25% acetonitrile in 0.1% formic acid, eluted samples were dried to completion in a SpeedVac at 60C, dissolved in 5 µL 0.1% formic acid, and stored at −20°C until MS analysis.

### Immunoprecipitation for MS analysis

For Dbn1 IP-MS analysis, PAS beads were bound to either IgG or the antibody raised against *Xenopus* Dbn1 (stock concentration 1 mg/mL) and at a beads-antibody ratio of 1:2 overnight at 4°C. Beads were then washed four times with ELB buffer and resuspended in IP buffer (ELB buffer supplemented with 0.35% NP-40, 5 mM NaF, 2 mM sodiumorthovanadate and 5 mM β-glycerophosphate). One volume of the egg extract reactions indicated was then added to 0.4 volumes of antibody-bound beads in a total of 5 volumes of IP buffer and incubated at 4°C for 1 hour with end-over-end rotation. Beads were washed thrice in ELB buffer supplemented with 0.25% NP-40 and 500 mM NaCl, transferred to LoBind tubes, and washed a further three times with ELB buffer supplemented with 500 mM NaCl. Immunoprecipitation of Cul1 and β-Trcp1 was performed with the indicated antibodies in essentially the same manner except, that one volume of unstimulated HSS was added to 0.67 volumes of beads in 3.67 volumes of ELB buffer and incubated 3h at 4°C, the washed thrice in ELB buffer supplemented with 0.25% NP-40, followed by three washes in ELB buffer. Beads were resuspended in 50 mM ammonium bicarbonate and samples subjected to on-bead digestion via addition of 100 ng modified sequencing grade Trypsin (Sigma) with incubations of 1 hour at 4°C and then overnight at 37°C with continuous mixing at 1200 rpm. Eluates were separated from beads by centrifugation through a 0.45 µM PVDF filter column (Millipore) and subjected to further 1 hour of in-solution digestion by addition of 100 ng additional Trypsin. Cysteines were subsequently reduced and alkylated by addition of 5 mM TCEP and 10 mM CAA for 30 min at 30°C and tryptic peptides were acidified and desalted on C18 StageTips as described above. Additionally, for the Dbn1 IP-MS experiment, samples were divided into two of which one half was desalted using low-pH clean-up as described above, while the other half was desalted using high-pH clean-up (Hendriks et al., 2018). High-pH clean-up was done essentially as described above except StageTips were equilibrated using 100 μl of methanol, 100 μl of 80 % ACN in 200 mM ammonium hydroxide, and two times 75 μl 50 mM ammonium. Samples were supplemented with 0.1 volumes of 200 mM ammonium hydroxide (pH >10), just prior to loading them on StageTip. The StageTips were subsequently washed twice with 150 μl 50 mM ammonium hydroxide, and afterwards eluted using 80 μl of 25 % ACN in 50 mM ammonium hydroxide.

### Whole proteome analysis

Volumes corresponding to 100 µg protein from three different batches of HSS extracts were diluted 100-fold in denaturing digestion buffer (6M Guanidine hydrochloride, 100 mM Tris, 5 mM TCEP, 10 mM CAA, pH 8.5), sonicated and digested using Lys-C (1:100 w/w; Wako) for 3 hours at RT. Digestions were subsdequently diluted with two volumes of 25 mM Tris pH 8.5 and further digested by addition of modified sequencing grade Trypsin (1:100 w/w) overnight at 37°C. Tryptic peptides were fractionated on-StageTip at high-pH essentially as described previously (Hendriks et al., 2018). Peptides were eluted from StageTips as eight fractions (F1-8) using 80 mL of 2, 4, 7, 10, 13, 17, 20, and 40% ACN in 50 mM ammonium. All fractions were dried to completion in LoBind tubes, using a SpeedVac for 3 h at 60°C, after which the dried peptides were dissolved using 12 µL of 0.1% formic acid.

### MS data acquisition

MS samples were analyzed on an EASY-nLC 1200 system (Thermo) coupled to either a Q Exactive HF-X Hybrid Quadrupole-Orbitrap mass spectrometer (Thermo) for the total proteome and DSB-UBIMAX samples, or an Orbitrap Exploris 480 mass spectrometer (Thermo) for the remaining MS experiments in this study. Separation of peptides was performed using 15-cm columns (75 mm internal diameter) packed in-house with ReproSil-Pur 120 C18-AQ 1.9 mm beads (Dr. Maisch). Elution of peptides from the column was achieved using a gradient ranging from buffer A (0.1% formic acid) to buffer B (80% acetonitrile in 0.1% formic acid), at a flow rate of 250 nL/min. For total proteome and DSB-UBIMAX samples, gradient length was 77 min per sample, including ramp-up and wash-out, with an analytical gradient of 55 minutes ranging from 5% to 25% buffer B for the total proteome and 52.5 minutes ranging from 10% to 25% buffer B for DSB-UBIMAX. For the remaining MS experiments, gradient length was 80 min per sample, including ramp-up and wash-out, with an analytical gradient of 57.5 minutes ranging in buffer B from 10% to 40% for DPC-UBIMAX samples and 52.5 minutes ranging in buffer B from 10% to 30% for IP-MS samples. The columns were heated to 40C using a column oven, and ionization was achieved using either a NanoSpray Flex ion source (Thermo) for the total proteome and DSB-UBIMAX, or a NanoSpray Flex NG ion source (Thermo) for the remaining MS experiments. Spray voltage set at 2 kV, ion transfer tube temperature to 275C, and RF funnel level to 40%. Samples were measured using 5 µL injections with different technical settings as detailed in the following. Measurements were performed with a full scan range of 300-1,750 m/z, MS1 resolution of 60,000, MS1 AGC target of 3,000,000, and MS1 maximum injection time of 60 ms for the total proteome and DSB-UBIMAX samples and MS1 resolution of 120,000, MS1 AGC target of ‘‘200’’ (2,000,000 charges), and MS1 maximum injection time to “Auto” for the remaining MS experiments. Precursors with charges 2-6 were selected for fragmentation using an isolation width of 1.3 m/z and fragmented using higher-energy collision disassociation (HCD) with a normalized collision energy of 28 for total proteome and DSB-UBIMAX and 25 for the remaining MS experiments. Precursors were excluded from resequencing by setting a dynamic exclusion of 45 s for total proteome and DSB-UBIMAX samples and 60 s with an exclusion mass tolerance of 20 ppm, exclusion of isotopes, and exclusion of alternate charge states for the same precursor for the remaining MS experiments. MS2 AGC target was set to 200,000 and minimum MS2 AGC target to 20,000 for total proteome and DSB-UBIMAX samples and MS2 AGC target to ‘‘200’’ (200,000 charges) with an MS2 intensity threshold of 230,000 or 360,000 for DPC-UBIMAX, Dbn1 IP-MS and β-Trcp1 IP-MS, respectively. For the total proteome samples, MS2 maximum injection time was 55 ms, MS2 resolution was 30,000, and loop count was 12. For DSB-UBIMAX samples, MS2 maximum injection time was 90 ms, MS2 resolution was 45,000, and loop count was 9. The MS2 settings were similar for the DPC-UBIMAX samples, except MS2 maximum injection time was set to “Auto”. This was also the case for the IP-MS samples, but while the Dbn1 IP-MS samples were aquired with MS2 resolution of 45,000 and a loop count of 9, the β-Trcp1 IP-MS samples were aquired with MS2 resolution of 15,000 and a loop count of 18. For the DPC-UBIMAX and IP-MS experiments, Monoisotopic Precursor Selection (MIPS) was enabled in ‘‘Peptide’’ mode.

### MS data analysis

All MS RAW data were analyzed using the freely available MaxQuant software (Cox and Mann, 2008), version 1.6.0.1. Default MaxQuant settings were used, with exceptions specified below. For generation of theoretical spectral libraries, the Xenopus laevis FASTA database was downloaded from Uniprot on the 13^th^ of May 2020 for the total proteome and UBIMAX experiments and on the 3^rd^ of September 2021 for the IP-MS experiments. In silico digestion of proteins to generate theoretical peptides was performed with trypsin, allowing up to 3 missed cleavages. Allowed variable modifications were oxidation of methionine (default), protein N-terminal acetylation (default) for all samples. For UBIMAX experiments, ubiquitylation of lysine and cysteine as well as carbamidomethyl on cysteine were additionally included as variable modifications. For IP-MS experiments, ubiquitylation of lysine and phosphorylation of serine and threonine were additionally allowed. Maximum variable modifications per peptide was reduced to 3. Label-free quantification (LFQ) (Cox et al., 2014) and iBAQ was enabled. For DSB-UBIMAX, LFQ was applied separately within parameter groups defined by sample type (controls *versus* ubiquitin target enriched samples). Stringent MaxQuant 1% FDR data filtering at the PSM and protein-levels was applied (default). Second peptide search was enabled. Matching between runs was enabled, with an alignment window of 20 min and a match time window of 1 min. For total proteome analysis, matching was only allowed within the same fractions and for IP-MS experiments within replicates of the same sample group. For the Dbn1 IP-MS experiment, dependent peptide search was additionally enabled.

### MS data annotation and quantification

The Xenopus laevis FASTA database downloaded from UniProt lacked comprehensive gene name annotation. Missing or uninformative gene names were, when possible, semi-automatically curated, as described previously (Gallina et al., 2021). Quantification of the MaxQuant output files (“proteinGroups.txt”) was performed using Perseus software (Tyanova et al., 2016) as was Pearson correlation and Principal Component analyses. For quantification purposes, all protein LFQ intensity values were log2 transformed, and filtered for presence in 4 of 4 replicates in at least one experimental condition for the DSB-UBIMAX experiment; 3 of 3 for total proteome, DPC-UBIMAX and β-Trcp1 IP-MS experiments and 4 of 8 in the Dbn1 IP-MS experiment. Missing values were imputed below the global experimental detection limit at a downshift of 1.8 and a randomized width of 0.3 (in log2 space; Perseus default). For the data presented in volcano- and scatter plots, statistical significance of differences was tested using two-tailed Student’s t-testing, with permutation-based FDR control applied at s0 values of 0.1 and proteins were filtered to be significantly enriched or depleted at FDR < 5%. Only proteins testing significantly enriched over both “no His” and “Ub E1i” controls for UBIMAX and over the mock control for the IP-MS experiments, respectively, with FDR < 5% when tested using one-tailed Student’s t-testing, with permutation-based FDR control applied at an s0 value of 0.1, were considered for further analysis. All tested differences, p-values, and FDR-adjusted q-values are reported in Supplementary tables S1, S3, S4 and S5. For hierarchical clustering analysis of ubiquitylated proteins robustly changing with DNA treatment only robustly changing proteins were considered for the analysis. Robustly changing was defined as proteins increased compared to the median in all four replicates of at least one sample group and decreased compared to the median in all four replicates of another sample group. Quantification of individual peptides or summed peptide abundances were derived from the MaxQuant output files (“evidence.txt”)

The original analysis of the total proteome included triplicate samples of both HSS and nucleoplasmic extract (NPE). However, as all *Xenopus* egg extract experiments in this study are otherwise performed in HSS, only this extract was included in the further analysis of the total proteome. The original analysis of the DPC-UBIMAX experiment included four replicates but one replicate was excluded due to significant technical variance. The samples of the Dbn1 IP-MS experiment was aquired as two technical replicates on the basis of C18 StageTip method (see above), with runs resulting from high-pH StageTip clean-up denoted by “H” in the raw files and replicates 01-04 in the analysis, while runs resulting from low-pH StageTip clean-up is denoted by “L” in the raw files and replicates 05-08 in the analysis. Furthermore, this experiment originally included a condition treated with ubiquitin E1 enzyme inhibitor and DSB-mimikcking plasmid DNA, but as this condition did not yield significant additional information, it was excluded for further analysis. Finally, in figure 3H, only the samples resulting from high-pH StageTip clean up are presented.

### Cell culture and ubiquitin pulldown

Human HeLa cell lines were cultured under standard conditions at 37°C and 5% CO_2_ in DMEM (Thermo) containing 10% FBS (v/v) and penicillin-streptomycin (Thermo). Cells were regularly tested negative for mycoplasma infection. To block Cullin E3 ligase activity, 1 µM neddylation E1 enzyme inhibitor (MLN4924, R&D systems) was added to the cell culture medium. To prevent ATM activity, the cell culture medium was supplemented with 10 µM ATM inhibitor (KU-55933, Selleckchem). Where indicated, cells were subjected to 10 Gy of ionizing radiation using a Smart X-ray machine (Yxlon).

For WB analysis, cells were lysed in RIPA buffer (140 mM NaCl, 10 mM Tris-HCl (pH 8.0), 0.1% sodium deoxycholate (w/v), 1% Triton X-100 (v/v), 0.1% SDS (w/v), 1 mM EDTA, 0.5 mM EGTA). For ubiquitin pulldown, cell lysates were prepared in lysis buffer (50mM Tris (pH 8.0), 1M NaCl, 5mM EDTA, 1% IGEPAL, 0.1% SDS). Lysates were sonicated once for 20 seconds with an amplitude of 75% on a hand held sonicator, before spinning down at full speed for 20 minutes on a 4°C centrifuge. Ubiquitin enrichment was performed using Halo-tagged MultiDsk TUBE (Wilson et al., 2012) preincubated with HaloLinkTM resin (G1913, Promega) for 1 hour with rotation at RT in binding buffer (100mM Tris (pH 7.5), 150mM NaCl, 0.05% IGEPAL). Excess protein was washed off with binding buffer supplemented with 1 mg/mL BSA. Cell lysates were added to the MultiDsk-bound resin and incubated with rotation overnight at 4°C. Samples were washed four times with MultiDsk lysis buffer before being eluted in 2x Laemmli sample buffer by boiling for 5 min at 95°C.

All lysis and wash buffers were supplemented with 1mM dithiothreitol (Sigma), complete EDTA-free protease inhibitor Cocktail Tablets (Roche), 1.25 mM N-ethylmaleimide (Sigma) and 50 µM PR-619 (Calbiochem).

### siRNA and plasmid transfections

The following siRNAs were used in this study to knock-down the expression of selected proteins: Non-targeting control (siCTRL) 5′-GGGAUACCUAGACGUUCUA-3′, siDBN1 5’-GGAGCUUUCGGGACACUUUtt −3’, siβ-Trcp1#1 5’-GUGGAAUUUGUGGAACAUCtt-3’, siβ-Trcp1#2 5’-AAGUGGAAUUUGUGGAACAUCtt-3’. All siRNAs were used at 20 nM concentrations and transfected with Lipofectamine RNAiMAX reagent (Thermo) according to the manufacturer’s instructions.

For transient overexpression of WT or S599A mutated DBN1, Full-length DBN1 (human) cDNA was inserted into pcDNA4/TO-EGFP through Gateway® cloning. The DBN1-S599A phospho-mutant was generated by Q5 mutagenesis with sgRNA-DBN1 5′-CCAGTGAGGGGTACTTCGCTCAATCACAGGAGG-3′. Plasmids were transfected with Lipofectamine 2000 (Thermo) according to the manufacturer’s instructions.

### *In silico* analysis of variant β-Trcp1 degron

A list containing the identity, motif sequence and sequence position of the human proteins containing a variant β-Trcp1 degron was generated by submitting the motif [DEST]-[DES]-G-x(2)-[ST]-Q to the ScanProsite tool ((de Castro et al., 2006) to scan against the UniProt Homo Sapiens (taxonomy ID: 9606) database. To this list was mapped the phosphorylation status and kinase relationship of the [ST]-Q site, if known, as retrived from the Phospho.ELM (Diella et al., 2008) and PhosphoSitePlus v6.7.1.1 (Hornbeck et al., 2015) databases.

### Quantification and statistical analysis

Bioinformatic analysis of mass spectrometry data were carried out with the Perseus software. Statistical significance of differences was tested using Student’s t testing, with permutation-based FDR-control applied at an s0 value of 0.1. Autoradiographs were quantified using ImageJ. Graphs and the statistical tests displayed in them were done in Prism (GraphPad) using the statistical tests indicated for each analysis. For all statistical analyses: **, p-value ≤ 0.01; ***, p-value ≤ 0.001; ****, p-value ≤ 0.0001. Error bars represent the standard error unless otherwise stated.

## Supplementary table legends

**Table S1. UBIMAX in response to DSBs.** MS analysis of ubiquitylated proteins enriched via the UBIMAX workflow (Figure 1A) from quadruplicate samples and according to the experimental outline in Figure 1D. Related to Figures 1D-I, 2D-I, S1D-I, and S2B. Proteins are enriched from reactions in the presence of “no His”, untagged recombinant ubiquitin and linearized plasmid DNA; “Ub E1i”, ubiquitin E1 inhibitor, recombinant His_6_-ubiquitin and linearized plasmid DNA; “no DNA”, recombinant His_6_-ubiquitin; “DNA”, recombinant His_6_-ubiquitin and undamaged plasmid DNA, “DSB”, recombinant His_6_-ubiquitin and linearized plasmid DNA.

**Table S2. Total egg extract proteome.** Whole proteome MS analysis of triplicate high-speed supernatant interphase egg extracts (HSS). Related to Figures 1F-G.

**Table S3. UBIMAX in response to DPCs.** MS analysis of ubiquitylated proteins enriched via the UBIMAX workflow (Figure 1A) from triplicate samples and according to the experimental outline in Figure 2A. Related to Figures 2A-I and S2B. Proteins are enriched from reactions in the presence of recombinant His_6_-ubiquitin and “no DNA”, buffer; “DNA”, undamaged plasmid DNA, “ssDNA-DPC”, plasmids carrying the M.HpaII protein crosslinked at a single-stranded DNA gap; “SSB-DPC”, plasmids carrying the Flp protein crosslinked at a single-strand break; “ssDNA-DPC + UbE1i”, ubiquitin E1 inhibitor and plasmids carrying the M.HpaII protein crosslinked at a single-stranded DNA gap; “SSB-DPC + UbE1i”, ubiquitin E1 inhibitor and plasmids carrying the Flp protein crosslinked at a single-strand break.

**Table S4. Dbn1 interactome in response to DSBs.** MS analysis of proteins and phosphorylation sites enriched via mock-or Dbn1 immunoprecipitation in quadruplicates and two technical replicates according to the experimental outline in Figure 3E. Related to Figures 3E-H, S3F, and S4A. Mock immunoprecipitations were performed in the presence of undamaged plasmid DNA, while proteins immunoprecipitated with Dbn1 antibodies were performed in the presence of “DNA”, undamaged plasmid DNA; “DSB”, linearized plasmid DNA”; “DSB + ATMi”, ATM inhibitor and linearized plasmid DNA; “DSB + MG262”, proteasome inhibitor and linearized plasmid DNA.

**Table S5. MS-based validation of β-Trcp1 antibodies.** MS analysis of proteins enriched via mock-, Cul1-, β-Trcp1-INT or β-Trcp1-NT immunoprecipitations from unstimulated egg extracts. Related to Figure S3G.

**Table S6. *In silico* analysis of a DDR-variant β-Trcp1 degron.** Proteome-wide sequence analysis of the occurrence of the [DEST]-[DES]-G-x(2)-[ST]-Q variant β-Trcp1 degron across all annotated human proteins. Phosphorylation status of the degron [ST]-Q site, according to the Phospho.ELM and PhosphoSitePlus (PSP) databases, is included.

## Acknowledgements

We thank members of the Nielsen and Duxin laboratories for feedback on the manuscript. We thank Philip Zegerman for the βTrcp1 encoding plasmid and Alex Bullock for the plasmids encoding Cul1-NT, Cul3a-NT and Cul5a-NT. The Novo Nordisk Foundation Center for Protein Research is supported financially by the Novo Nordisk Foundation (grant agreement NNF14CC0001). The work carried out in this study was in part supported by the Novo Nordisk Foundation Center for Protein Research, the Novo Nordisk Foundation (NNF14CC0001, NNF13OC0006477; M.L.N., and NNF21OC0071976; J.P.D.), The Danish Council of Independent Research (8020-00220B, M.L.N.), The Danish Cancer Society (R146-A9159-16-S2, M.L.N.). The proteomics technology applied was part of a project that has received funding from the European Union’s Horizon 2020 research and innovation program under grant agreement EPIC-XS-823839 (M.L.N.). J.A.G. was funded by the European Union’s Horizon 2020 research and innovation program (Marie-Skłodowska-Curie grant agreement no. 860517 (UBIMOTIF)). This project has also received funding from the Medical Research Council (MR/K007106/1, A.G.). Figures 4A, 4D and 4G were created using BioRender.com.

## Author contributions

C.S.C., J.P.D., M.L.N. conceived the project. C.S.C., I.A.H., M.L.N. designed the MS experiments and UBIMAX method. C.S.C. performed the MS experiments. C.S.C., I.A.H. analyzed the MS data. C.S.C., I.A.H., J.P.D., M.L.N. interpreted the MS data. C.S.C., E.S.K., J.P.D. designed the Xenopus egg extract experiments. C.A., A.G. produced the recombinant dominant negative Cullin proteins. Z.F. generated the SSB-DPC plasmid DNA substrate. C.S.C., E.S.K. performed the Xenopus egg extract experiments. C.S.C., E.S.K., J.P.D., M.L.N. interpreted the Xenopus egg extract experiments. C.S.C., E.S.K., J.A.G., N.M., J.P.D. designed the human cell experiments. J.A.G. performed the human cell experiments. C.S.C., E.S.K., J.A.G., N.M., J.P.D., M.L.N. interpreted the human cell experiments. C.S.C. performed the *in silico* analysis of the variant βTRCP1 degron. C.S.C. prepared the figures and wrote the manuscript with feedback from J.P.D. and M.L.N. E.S.K. created the models of Figures 4A, 4D and 4G. All authors provided critical review of the manuscript.

---

**Figure S1.**
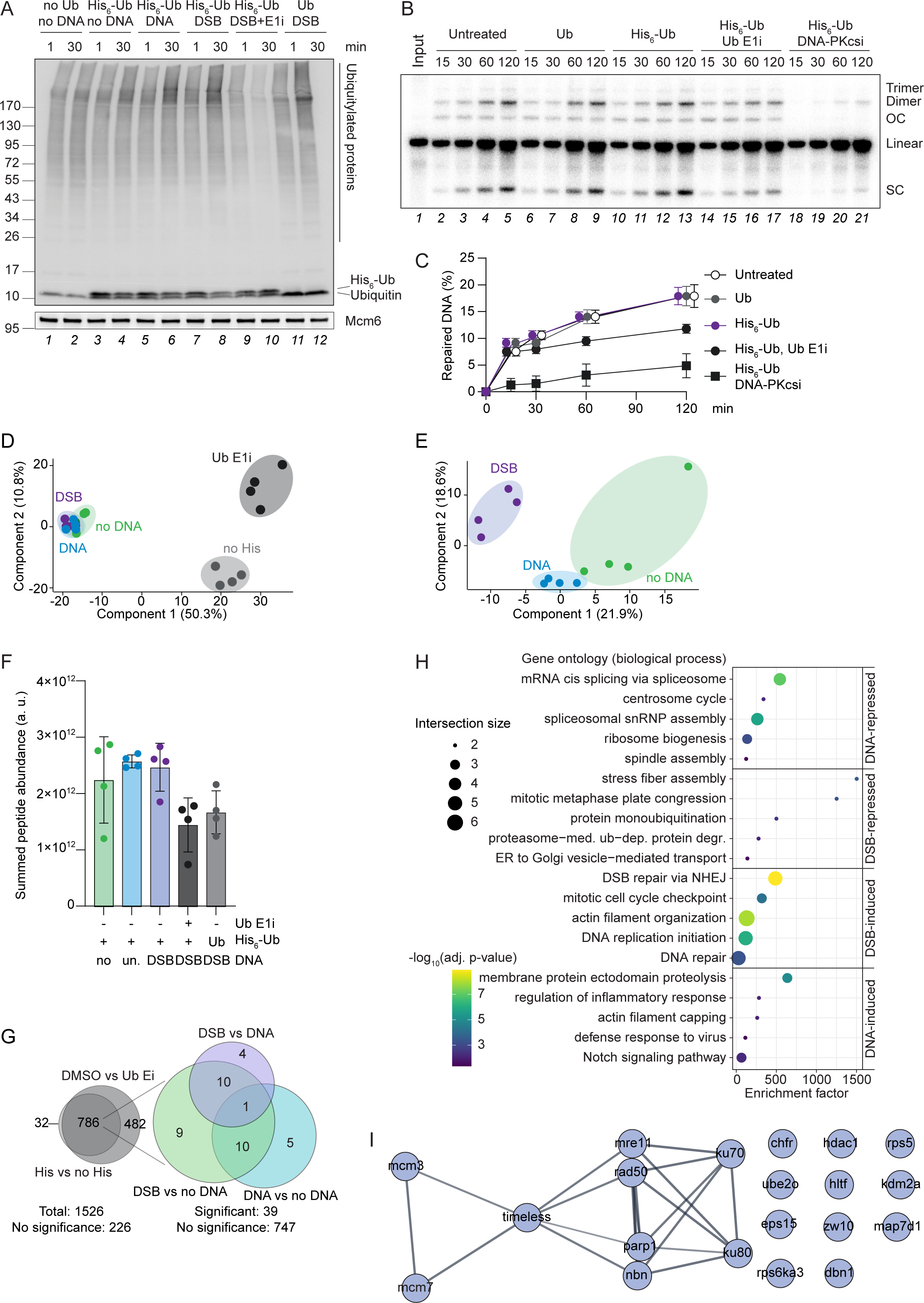
UBIMAX efficiently and specifically detects ubiquitin-conjugated proteins in response to DSBs. Related to Figure 1. **A.** WB analysis of ubiquitin from the experiment shown in Figure 1C. Mcm6 served as a loading control. **B.** Egg extracts were left untreated or supplemented with ubiquitin E1 inhibitor (“Ub E1i”) or DNA-PKcs inhibitor (“DNA-PKcsi”) prior to addition of recombinant untagged ubiquitin (“Ub”) or His_6_-Ubiquitin (“His_6_-Ub”) as indicated. Reactions were initiated by addition of radioactively labelled, linearized plasmid DNA. Samples were collected at the indicated timepoints, the DNA recovered, and analysed by agarose gel electrophoresis. Radioactively labelled, linearized plasmid DNA alone served as an input control. A representative experiment is shown. SC; supercoiled, OC; open circular. **C.** Quantification of the experiment shown in (B). Within each time point, the signal from radioactively labelled repair products (supercoiled, dimers and trimers) are summarised and quantified as percent of total radioactive signal within each sample and normalized to the 0-minute time point. Error bars represent standard error of the mean. N=2-4. **D-E.** Principal component analysis of all sample groups (**D**) and ubiquitin target enriched sample groups only (**E**). The analysis is based on proteins that are detected in all four replicates in at least one sample group. The principal components displayed represent the greatest degree of variability observed within the data. Eigenvalues are displayed on the axes. **F.** Mean summed peptide abundance of four replicates across all sample groups. Error bars represent standard deviations. no, no DNA (buffer); un., undamaged plasmid DNA; DSB, linearized plasmid DNA; a.u., arbitrary units. **G.** Proportionally scaled Venn diagrams showing the overlap of proteins significantly enriched over controls either in the presence *versus* absence of His_6_-ubiquitin, (“His vs no His”) or in the absence *versus* presence of ubiquitin E1 inhibitor (“DMSO vs Ub E1i”) (*left*) and the subset of proteins significantly enriched over controls whose enrichment further changed significantly upon DNA treatment – either in the presence *versus* absence of undamaged plasmid DNA (“DNA vs no DNA”) or linearized plasmid DNA (“DSB vs no DNA”) or with undamaged *versus* linearized plasmid DNA (“DSB vs DNA”) (*right)*. Isoforms were excluded if annotated with a gene name. Significant enrichment of the proteins populating the diagrams were determined by one- and two-tailed Student’s *t* testing, for left and right Venn diagrams respectively, with S0 = 0.1 and FDR ≤ 0.05. **H.** Enrichment analysis of Gene Ontology biological processes significantly represented (FDR ≤ 0.05) in the clusters of the hierarchical clustering analysis shown in Figure 1H, when compared to a total *Xenopus laevis* proteome derived from UniProt. Significance was determined via two-tailed Fisher’s Exact testing with Benjamini–Hochberg correction for multiple hypotheses testing. Selected categories are shown. Complete list is provided in Table S1. **I.** STRING network analysis of the proteins contained in the cluster labelled “DSB-induced” in Figure 1G. Proteins were queried against the STRING *Xenopus* (Silurana) *tropicalis* database at a confidence level of 0.7.

**Figure S2.**
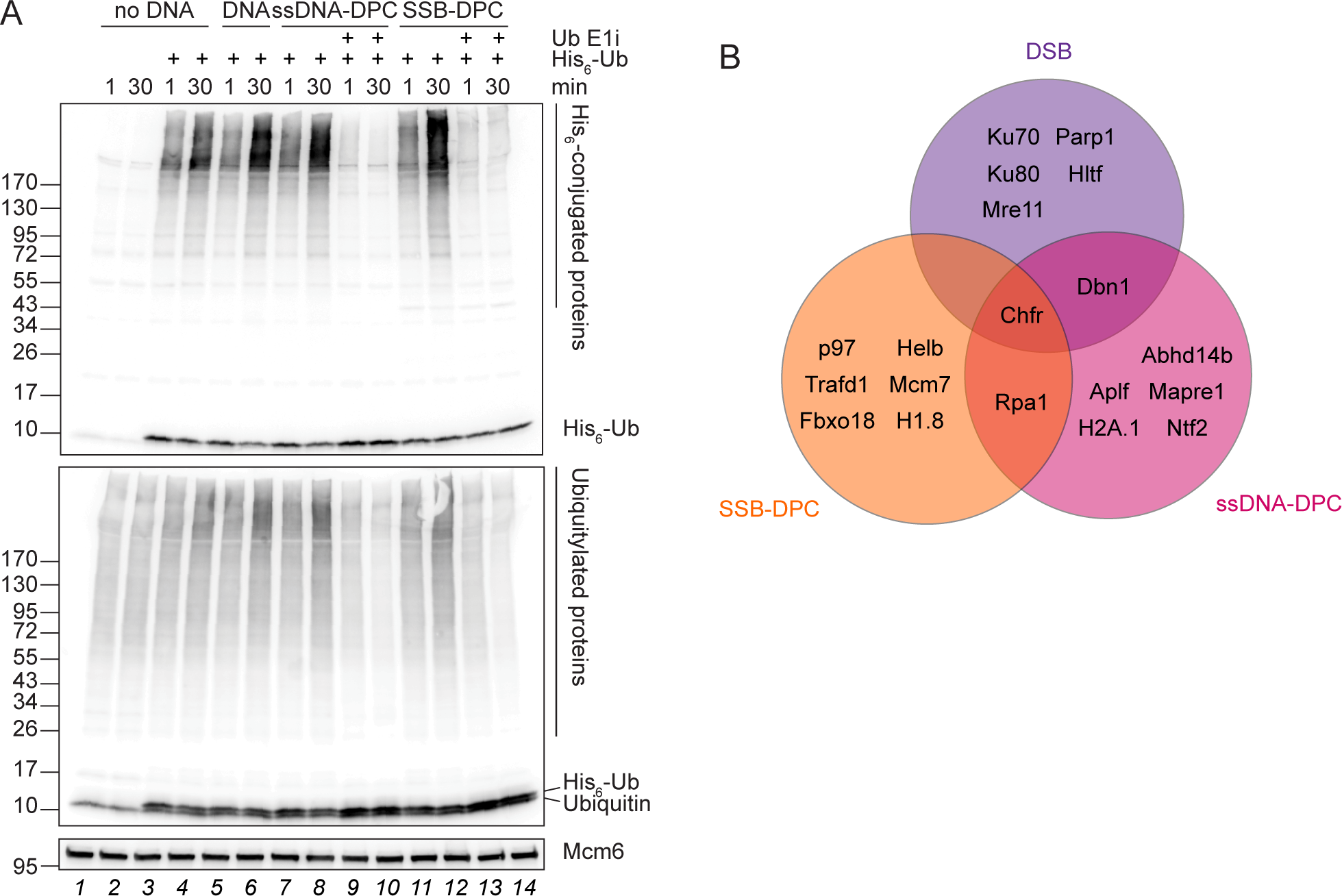
UBIMAX identifies DNA damage specific ubiquitylation events. Related to Figure 2. **A.** Egg extract reactions were performed as described for Figure 2A, except reactions were performed either in the absence or presence of His_6_-ubiquitin as indicated. Samples were transferred to sample buffer at 1 or 30 min after initiation and analysed by WB using antibodies against the His-tag and ubiquitin. Mcm6 served as a loading control. **B.** Venn diagram showing the overlap of ubiquitylated proteins detected by UBIMAX as significantly enriched in response to linearized plasmid DNA (“DSB”), plasmids carrying the M.HpaII protein crosslinked at a single-stranded DNA gap (“ssDNA-DPC”), or plasmids carrying the Flp protein crosslinked at a single-strand break (“SSB-DPC”), all compared to undamaged plasmid DNA. Only proteins significantly enriched, as determined by two-tailed Student’s *t* testing, with S0 = 0.1 and FDR ≤ 0.01, are shown.

**Figure S3.**
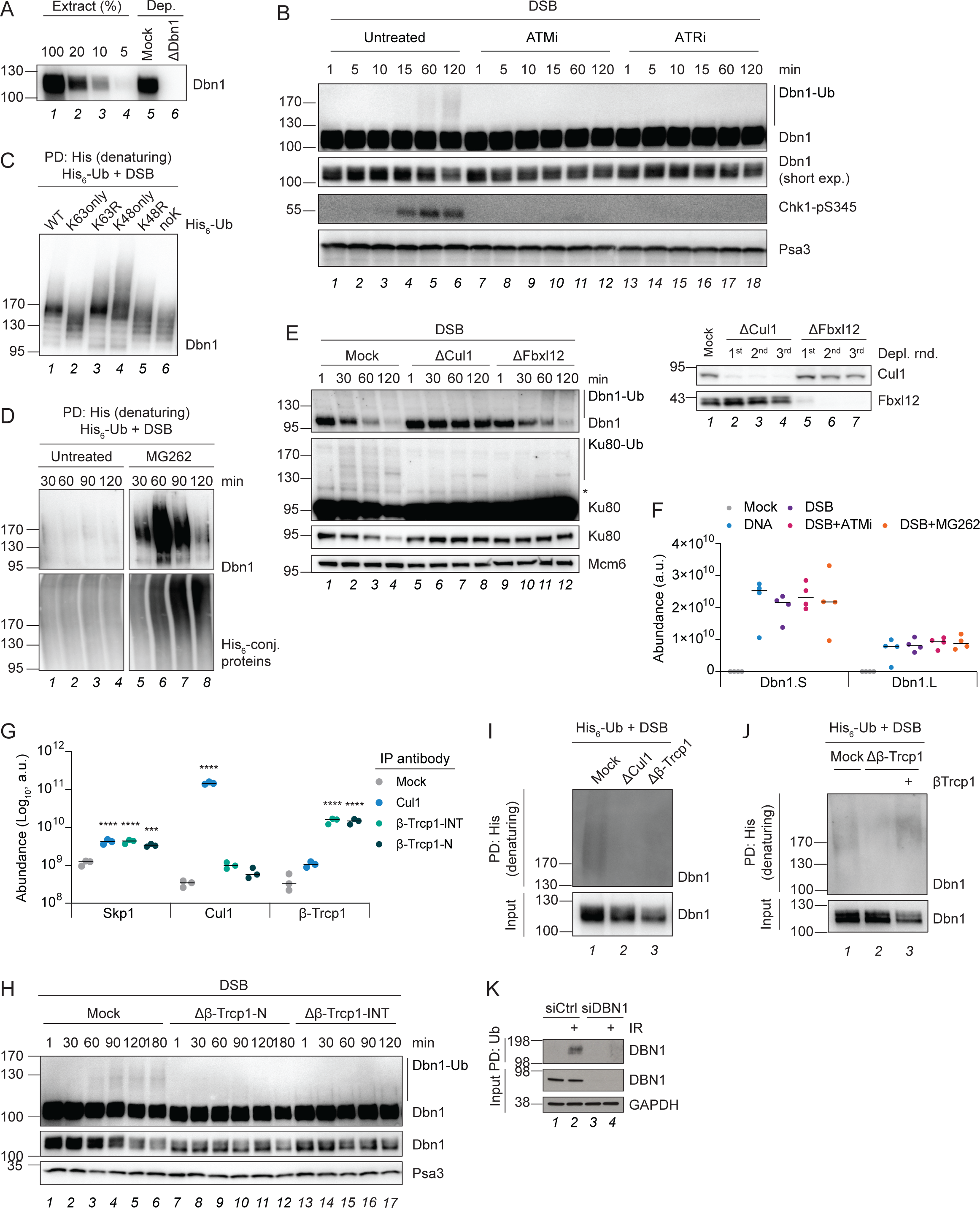
DDR-dependent and SCF^β-Trcp1^-mediated K48-linked ubiquitylation of Dbn1 results in proteasomal degradation. Related to Figure 3. **A.** Dbn1 immunodepletion control of the experiment shown in Figure 3A. An egg extract dilution range is shown for comparison. **B.** Egg extracts were left untreated or supplemented with ATM or ATR inhibitor (“ATMi”, “ATRi”) prior to addition of linearized plasmid DNA (“DSB”). Samples were transferred to sample buffer at the indicated timepoints and analysed by WB using antibodies against Dbn1 and Chk1-pS345. Orc2 served as a loading control. **C.** His_6_-Ubiquitin (“His_6_-Ub”) WT or the indicated mutants were added to egg extracts at a final concentration of 1 µg/µl. Reactions were initiated by addition of linearized plasmid DNA, samples collected after 30 minutes, and subjected to denaturing His-ubiquitin pulldown followed by WB analysis using antibodies against Dbn1. PD, pulldown. **D.** Egg extracts were left untreated or supplemented with proteasome inhibitor (MG262) and His_6_-ubiquitin prior to addition of linearized plasmid DNA. Samples were collected at the indicated timepoints and subjected to denaturing His-ubiquitin pulldown followed by WB analysis using antibodies against Dbn1. Ubiquitin served as a pulldown control. **E.** Linearized plasmid DNA was added to Mock-, Cul1-or Fbxl12-immunodepleted egg extracts and samples transferred to sample buffer at the indicated timepoints. Samples were analysed by WB using antibodies against Dbn1 and Ku80 (long and short exposures shown). Mcm6 served as a loading control. Cul1-and Fbxl12 immunodepletion controls are shown to the right. Depl. rnd, immunodepletion round. * denotes an unspecific band. **F.** Abundance distribution of the two *Xenopus laevis* Dbn1 isoforms, Dbn1.S and Dbn1.L, across the sample groups of the Dbn1 IP-MS experiment outlined in Figure 3E. Horizontal lines indicate the median. N=4. a.u., arbitrary units. **G.** Unstimulated egg extracts were subjected to immunoprecipitation using IgG-(“mock”), Cul1-, β-Trcp1-INT, or β-Trcp1-NT antibodies, followed by MS analysis. Shown are the abundance distributions of Skp1, Cul1, and β-Trcp1. Horizontal lines indicate the median and significance was determined by one-way ANOVA with Dunnett’s multiple comparisons test for all immunoprecipitation conditions compared to the mock control. N=3. a.u., arbitrary units. **H.** Linearized plasmid DNA was added to mock-or β-Trcp1-immunodepleted egg extracts (using either β-Trcp1-INT or β-Trcp1-NT antibodies) and samples transferred to sample buffer at the indicated timepoints. Samples were analysed by WB using antibodies against Dbn1 (long and short exposures shown). Mcm6 served as a loading control. **I.** His_6_-ubiquitin and linearized plasmid DNA were added to mock-, Cul1-, or β-Trcp1-immunodepleted egg extracts. Samples were collected from immunodepleted egg extract prior to addition of DNA (“input”), and 60 minutes after addition of linearized DNA for denaturing His-ubiquitin pulldown. Samples were analysed by WB using antibodies against Dbn1. **J.** Recombinant β-Trcp1 protein or buffer was added to mock- and β-Trcp1-immunodepleted egg extracts as indicated prior to addition of His_6_-ubiquitin and linearized plasmid DNA. Input and pulldown samples were collected 60 minutes after addition of linearized plasmid DNA and processed as described in (J). **K.** HeLa cells were transfected with control or siRNA targeting DBN1 before being subjected or not to 10 Gy ionizing radiation (IR). Lysates were harvested 30 minutes after irradiation, subjected to ubiquitin pulldown, and analysed along with whole cell extracts (“input”) by WB using antibodies against DBN1. GAPDH served as a loading control. PD, pulldown; Ub, ubiquitin.

**Figure S4.**
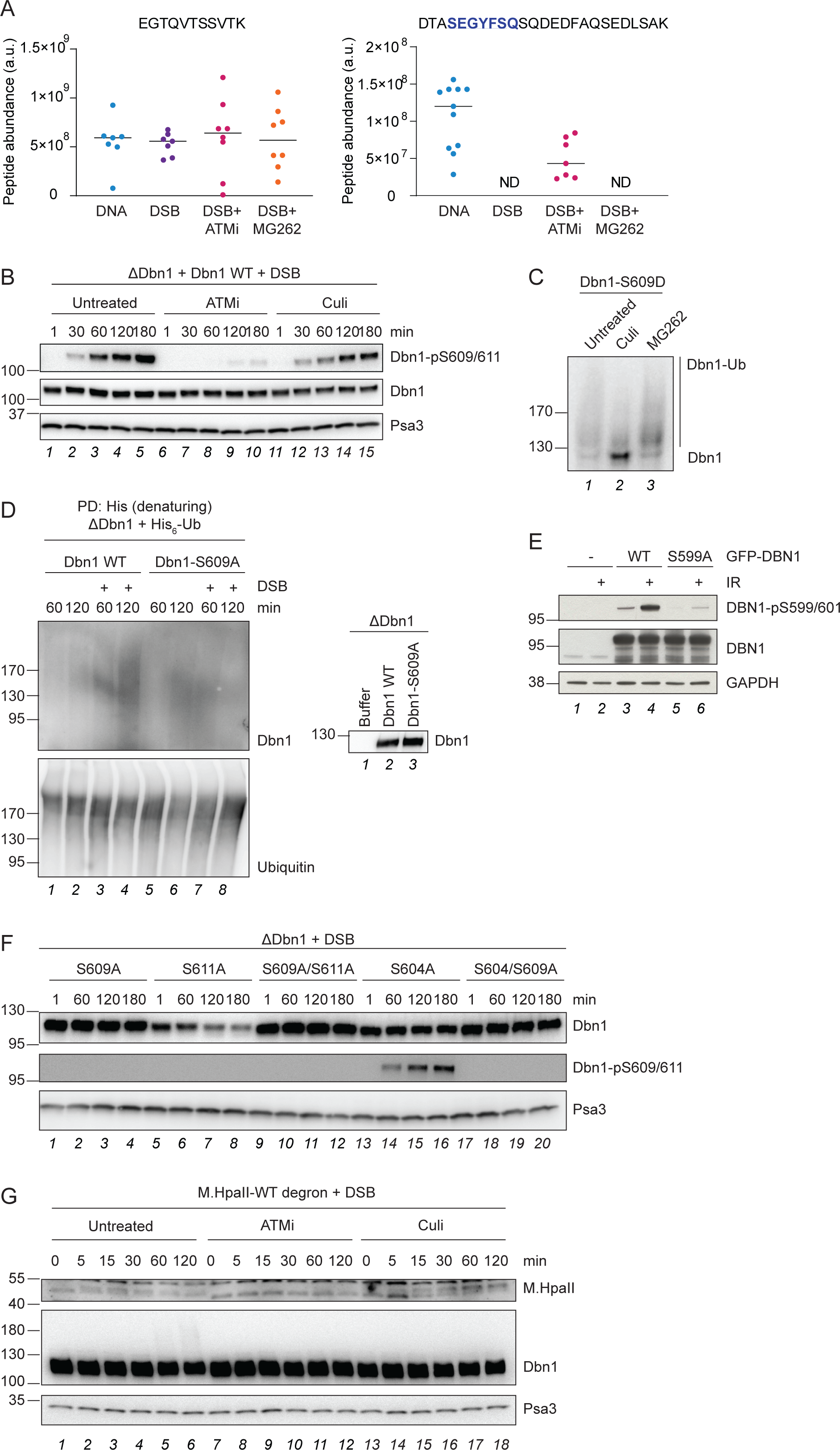
A variant β-Trcp1 degron is necessary and sufficient for inducing Dbn1 and general protein degradation in response to DSBs. Related to figure 5. **A.** Abundance of the indicated tryptic Dbn1 peptides across the samples of the Dbn1 IP-MS experiment outlined in Figure 3E. Horizontal lines indicate the median. a.u., arbitrary units. **B.** Dbn1-immunodepleted egg extracts were left untreated or supplemented with ATM inhibitor (“ATMi”) or neddylation E1 inhibitor (“Culi”), prior to addition of recombinant Dbn1 protein and linearized plasmid DNA (“DSB”). Samples were transferred to sample buffer at the indicated timepoints and analysed by WB using antibodies against Dbn1-pS609/S611 and total Dbn1. Psa3 served as a loading control. **C.** Rabbit reticulocyte lysates (*in vitro* translation system, IVTS) were left untreated or supplemented with neddylation E1 inhibitor or proteasome inhibitor (MG262) before initiating protein translation by addition of a plasmid encoding Dbn1-S609D. Samples were retrieved after 90 minutes and analysed by WB using antibodies against Dbn1. **D.** Recombinant Dbn1 protein, either WT or S609A mutant, was added to Dbn1-immunodepleted egg extracts prior to addition of 1 µg/µl His_6_-ubiquitin and undamaged-or linearized plasmid DNA (“DSB”). Samples were collected prior to addition of linearized plasmid DNA for Dbn1-immunodepletion and protein add-back controls (*right*), and 60 minutes after addition for denaturing His-ubiquitin pulldown (*left*). Samples were analysed by WB using antibodies against Dbn1. Ubiquitin served as a pulldown control. **E.** HeLa cells were transiently transfected with control plasmid (“-”) or plasmids encoding GFP-DBN1 (“WT”) or GFP-DBN1-S599A (“S599A”) for 24 hours before being subjected or not to 10 Gy IR. Lysates were harvested 30 minutes after irradiation and analysed by WB using antibodies against phosphorylated DBN1 (DBN1-pS599/601) and total DBN1. GAPDH served as a loading control. **F.** The indicated recombinant Dbn1 mutants were added to Dbn1-immunodepleted egg extracts prior to addition of linearized plasmid DNA. Samples were collected at the indicated timepoints and analysed by WB using antibodies against total and phosphorylated Dbn1. Psa3 served as a loading control. **G.** Recombinant M.HpaII protein tagged with the variant β-Trcp1 degron was mixed with egg extracts before supplementing or not with ATM-or neddylation E1 inhibitor. Reactions were initiated by addition of linearized plasmid DNA, samples were transferred to sample buffer at the indicated timepoints and analysed by WB using antibodies against M.HpaII and Dbn1. Psa3 served as a loading control.

